# Myelin basic protein is an RNA chaperone in microglial nuclear retro-transport

**DOI:** 10.64898/2026.05.14.725089

**Authors:** Guy Lam, Zhao Yang Xu, Adrien M Vaquié, Stavros Vagionitis, Michael Perry, Omar de Faria, Georgios Solomou, John H Stockley, Gemma C Girdler, Daniel Yamamoto, Juan A Oses, Qianqian Zhang, Gregory Jordan, Laura R Morcom, Jacob Stillman, Hani Mousa, Alma Burlingame, Murray Stewart, Hauke B. Werner, András Lakatos, Harry Bulstrode, Dorothy P. Schafer, Joanne Jones, Ragnhildur T Karadottir, David H Rowitch

**Affiliations:** Cambridge Stem Cell Institute, University of Cambridge, Puddicombe Way, Cambridge Biomedical Campus, Cambridge CB2 0AW, UK; Department of Paediatrics, University of Cambridge, Cambridge CB2 0QQ, UK; Departments of Pediatrics and Neurosurgery, Cedars-Sinai Guerin Children’s, Los Angeles, CA 90048, USA; Department of Veterinary Medicine, University of Cambridge, Cambridge CB3 0ES, UK; Division of Academic Neurosurgery, Department of Clinical Neurosciences, University of Cambridge, Cambridge CB2 0QQ, UK; Department of Pharmaceutical Chemistry, University of California, San Francisco, San Francisco, CA 94158, USA; Department of Neurobiology, Brudnick Neuropsychiatric Research Institute, University of Massachusetts Chan Medical School, Worcester, MA 01605, USA; Department of Clinical Neurosciences, University of Cambridge, Cambridge CB2 0QQ, UK; MRC Laboratory of Molecular Biology, Francis Crick Avenue, Cambridge CB2 0QH, UK; Neurochemistry Group, Max Planck Institute for Multidisciplinary Sciences, Göttingen 37077, Germany; Faculty of Biology and Psychology, University of Göttingen, Göttingen 37073, Germany

**Keywords:** Myelin Basic Protein, Nuclear pore transport, mRNA, Myelin, Ivermectin, Importin, Karyopherin, Microglia, gene editing, Multiple Sclerosis, neuroinflammation

## Abstract

CNS oligodendrocytes generate myelin, an RNA-containing proteolipid substance that enhances axonal transmission. In multiple sclerosis (MS), myelin debris is phagocytosed by microglia (MG), and prior studies have detected myelin-derived mRNA in MG nuclei, suggesting a retrograde transport pathway. We report myelin basic protein (MBP) is a nucleic acid–binding and trafficking protein. We found that retro-transport of myelin RNA into the MG nucleus was phagocytosis and importin-dependent. Transcriptomic and proteomic analyses of MG nuclei revealed enrichment of myelin mRNAs and proteins, with MBP singularly detected in soluble and chromatin-associated fractions. MBP bound mRNA with high affinity (Kd ≈ 0.30 nM) and was sufficient to facilitate MG RNA nuclear import *in vitro* and *in vivo*. Functionally, MBP mediated the delivery of small interfering RNAs for targeted knockdown of *toll-like receptor 4*. These findings indicate MBP as an RNA-binding protein capable of MG nuclear import, providing insight into neuroinflammatory pathology of MS.

## Introduction

Myelinating oligodendrocytes of the central nervous system (CNS) first appeared during evolution in jawed fishes ^1–3^. CNS myelination enhances conduction velocity^3^, maintains axon nutrition and integrity^4^, protects axons against inflammatory and oxidative insults^5^, modulates neural circuitry plasticity^6,7^ and is associated with adaptive task-learning ^8^.Oligodendrocyte precursor cells (OPCs) dramatically expand their surface area (∼6500-fold)^9,10^ during maturation to oligodendrocytes^11–15^, a biosynthetic process that requires tight coordination of myelin protein expression^16,17^, RNA localization^18–20^, and cytoskeletal remodelling ^21–24^ to generate compact myelin^25^ and specialized non-compact regions^20,26^. Non-compact myelin regions retain cytoplasm enriched in microtubules^27,28^, ribosomes/polysomes^29^, and vesicular transport machinery^12,28^, providing a local reservoir for translation of key myelin proteins^16,18–20,29–31^ that supports white matter integrity^32,33^ and function^29,30^. Functional integrity of myelin depends on myelin basic protein (MBP), which is essential for myelin sheath compaction^25,35^, involving interactions with myelin lipids ^25,36–40^. MBP is intrinsically disordered^41–43^ and conformationally flexible, enabling binding to diverse partners ^35,41,43,44^. The 21.5-kDa MBP isoform contains a non-traditional proline-tyrosine (PY)-type nuclear localization signal^45^, suggesting that specific MBP isoforms are karyophilic^46,47^, and may regulate transcriptional programs during oligodendrocyte proliferation ^47^ and disease states ^48^.

Multiple Sclerosis (MS) is a leading cause of neurological disability in young adults ^49,50^, with over 2.8 million individuals affected worldwide. In MS, adaptive immune system-mediated damage to myelin and oligodendrocytes leads to myelin destruction ^50,51^. Remyelination can occur early in the disease, leading to a relapsing–remitting clinical course, with later transition to progressive disability^49,52^. MS outcomes have been substantially improved by disease-modifying therapies that suppress disease activity and reduce relapses predominantly through modulation or depletion of peripheral adaptive immune cells, including both B- and T-cell compartments ^49,53–56^. While immune system myelin pruning occurs in development^57,58^, adaptive immune-mediated myelin damage generates debris that is avidly phagocytosed by resident microglia and infiltrating macrophages (MG/MP)^59^. Interestingly, several single-nucleus RNA sequencing studies in MS^60–66^ and Alzheimer’s disease models^67^ have revealed myelin mRNAs in MG nuclei, suggesting a retrograde RNA nuclear transport mechanism in neuropathological conditions ^60,61,67^.

As illustrated in **Figure 1(e)**, microglial uptake of myelin debris is mediated by the fatty acid translocase receptor CD36^68,69^ and NRF2 signalling ^70^ and can be blocked by the inhibitor sulfo-N-succinimidyl oleate (SSO) ^68,69^. Subsequent phagocytic processing depends on microtubule-driven cytoskeletal dynamics ^71,72^, vesicle trafficking, and phagosome maturation ^72,73^, that are sensitive to colchicine ^74,75^. In mammalian somatic cells, mRNA trafficking is generally unidirectional from the nucleus to the cytoplasm following transcription and processing ^76^, whereas retrograde nuclear import of single strand RNA is a strategy exploited by influenza^77,78^. This relies on viral chaperone proteins bearing nuclear localization signals (NLS)^79^ that engage karyopherins/importins^80^ to drive active nuclear import through the nuclear pore complex under Ran-GTP control ^79–83^; ivermectin^84–86^ and wheat germ agglutinin ^87,88^ can disrupt importin-dependent transport, highlighting pharmacologic checkpoints in this pathway.

**Figure 1.**
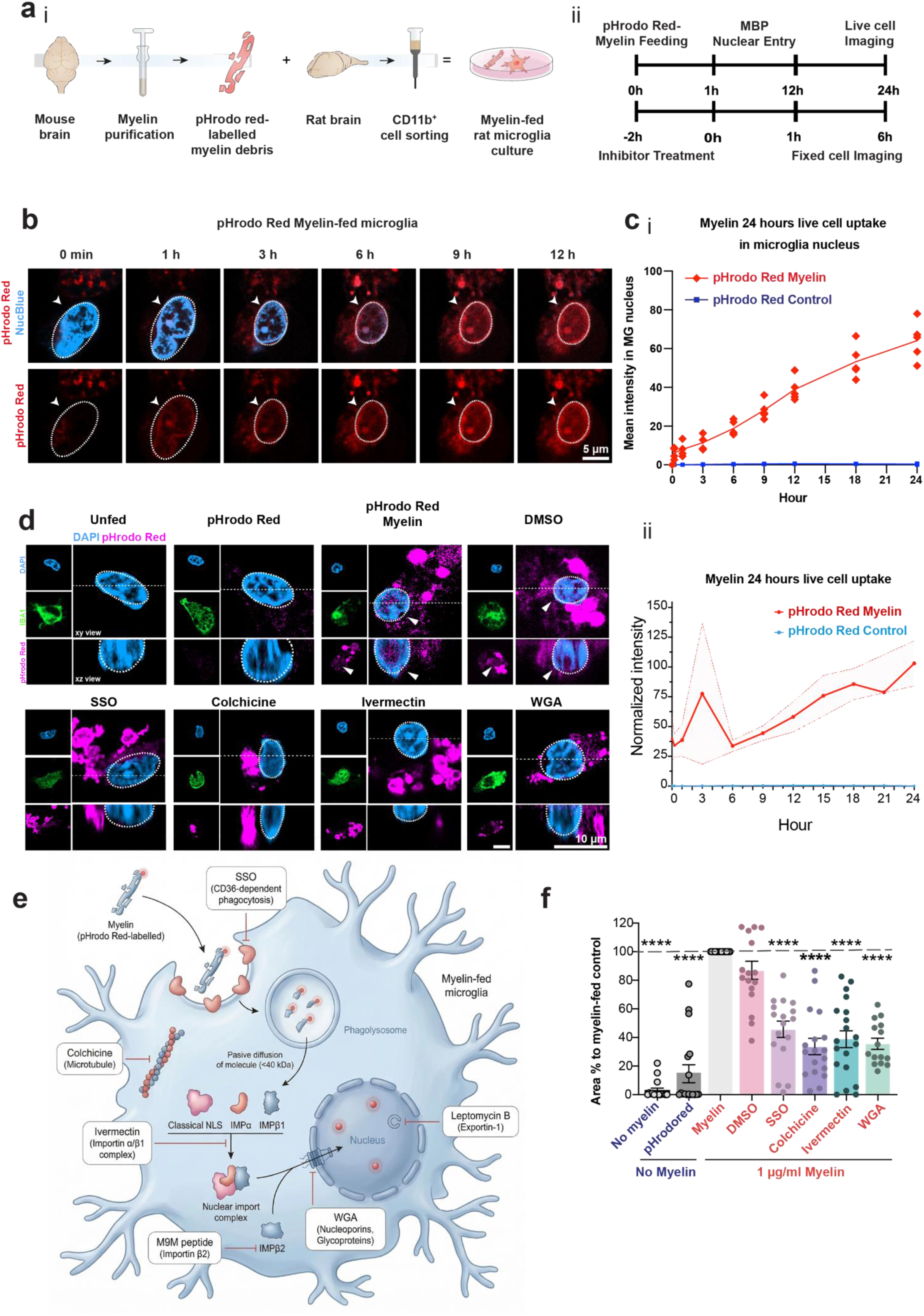
Myelin proteins access the microglial nucleus via Importins. **a,i,** Schematic of myelin purification from mouse brain, pHrodo Red labelling of myelin debris, isolation of CD11b^+^ microglia from rat brain, and establishment of myelin-fed microglia cultures. **a,ii**, Timelines for live-cell (b, c) and fixed-cell (d, f) imaging shown, including inhibitor pre-treatment where indicated. b, Representative live-cell confocal images of microglia (MG) incubation with pHrodo Red–labelled myelin over time (0–12h). Fluorescence progressively accumulates within the nucleus (outlined; NucBlue Live Cell nuclear stain), first observable at 1 h after myelin addition at 0 min and increasing thereafter. By 12 h, nuclear fluorescence is prominent. Arrowheads indicate nuclear enrichment. Scale bar, 5 μm. **c,i**, Quantification plot of b. Data are presented as mean fluorescence intensity over time (h) in the microglial nucleus. Each data point represents an individual cell. c,ii, Quantification plot of combined live-cell uptake experiments. Normalized mean fluorescence intensity over time (h) in the microglial nucleus over 24 h (mean ± s.e.m). Data normalized from three biological repeats (myelin preparations). Statistical analysis was performed using two-way ANOVA followed by Dunnett’s multiple comparisons test. \*\*\*\**p <* 0.0001, for myelin-fed versus pHrodo red-fed control. **d,** Representative confocal images of microglial cells, treated with pHrodo Red-labelled myelin (magenta) in the presence of the indicated inhibitors or DMSO control, and fixed at 6 h. Single optical sections are shown. Individual channels are displayed to the left for each condition, and the lower panels show XZ sections along the dotted line. pHrodo Red–labelled myelin is shown in magenta, IBA1 (microglial marker) in green and DAPI (nuclei) in blue. Conditions (left to right): (top) unfed; pHrodo Red control (1 μg ml⁻¹); pHrodo Red–labelled myelin (1 μg ml⁻¹); myelin + DMSO (1%); (bottom) myelin + SSO; myelin + colchicine; myelin + ivermectin; myelin + wheat germ agglutinin. Nuclear localization of pHrodo Red–labelled myelin is observed in myelin-fed cells and is reduced by inhibition of CD36 (SSO), microtubules (colchicine), importin α/β1 (ivermectin) or nucleoporins (wheat germ agglutinin). Scale bars, 10 μm. **e**, Schematic for myelin uptake and nuclear entry in microglia. pHrodo Red–labelled myelin is internalized via CD36-dependent phagocytosis, trafficked through the endo–lysosomal system, and subsequently engages importin-dependent nuclear import. Pharmacological inhibitors target distinct steps: sulfo-N-succinimidyl oleate (SSO; CD36-dependent uptake, 25 μM), colchicine (microtubules, 0.2 μg ml⁻¹), ivermectin (importin α/β1 complex, 1 μM), M9M peptide (importin β2, 10 μM), wheat germ agglutinin (WGA; nucleoporins, 100 μg ml⁻¹), and leptomycin B (exportin-1, 10 nM). **f**, Quantification plot of d. Nuclear pHrodo red^+^ area/ object (pHrodo red^+^ Iba1^+^ DAPI^+^ area in μm^2^/ pHrodo red^+^ microglial objects), normalized to myelin-fed positive control). Data are presented as mean ± s.e.m. (*n =* 19 independent biological replicates, each representing a myelin preparation and microglia isolation). Each data point corresponds to one biological replicate, calculated as the normalized value averaged across >600 cells. Statistical significance was assessed using one-way ANOVA followed by Dunnett’s multiple-comparisons test. Normality and homogeneity of variance were assessed using Shapiro–Wilk and Brown–Forsythe tests, respectively. \*\*\*\**p <* 0.0001 versus myelin-fed control.

Here we investigated the mechanism by which myelin debris mRNA is retrotransported into the MG nucleus. We show that myelin proteins are rapidly assimilated into the rat MG cytoplasm and nucleus after myelin feeding in culture. We found selective uptake of myelin-associated mRNAs and proteins, particularly MBP, that was phagocytosis- and importin-dependent. We show that MBP is a non-selective mRNA-binding protein, facilitating retrograde transport of RNA cargo to the MG nucleus *in vitro* and *in vivo*. Interestingly, MBP facilitates delivery of siRNA for functional knockdown of *Toll-like receptor 4 (TLR4)*, suggesting potential for MBP – nucleic acid complexes to regulate MG responses.

## Results

### Nuclear uptake of myelin proteins is phagocytosis and nuclear pore complex-dependent

To investigate the kinetics of myelin protein import into the microglial (MG) nucleus, we fed myelin biochemically purified from mouse brains (Extended Data **Fig. 1**) to rat primary MG cultures. Visualization of myelin protein trafficking was achieved by pHrodo Red labelling (**Fig. 1ai**) and we conducted live cell imaging for 24 hrs post feeding (**Fig. 1aii**). We observed pHrodo Red signal in the nucleus as early as 1 hour (**Fig. 1b**), and the intensity continued to increase over the time course (**Fig. 1c**; two-way ANOVA, *p =* 4.45 × 10⁻⁶). In contrast, control cultures treated only with pHrodo Red showed no significant accumulation of signal in the nucleus (Extended Data **Fig. 2**, **Suppl. Video 1**). We noted that within 1 hr of exposure to myelin, MG adopted an amoeboid-like morphology (**Suppl. Video 2**), reminiscent of that observed in experimental demyelination models^89,90^ and human MS ^91,92^.

To test if the nuclear import mechanism was phagocytosis-, cytoskeletal transport-and/or nuclear pore complex-dependent, we treated rat MG with several inhibitors (illustrated in **Fig. 1e**) 2 hrs before myelin feeding, then imaged 6 hrs after feeding (**Fig. 1aii**). As shown (**Fig. 1d, f**), we identified significant reduction in the nuclear pHrodo Red^+^ area/ object (pHrodo Red^+^ Iba1^+^ DAPI^+^ area in μm^2^/ pHrodo Red^+^ MG cell objects averaged to positive control) in response to several inhibitors. Regarding phagocytosis, incubation with the CD36 inhibitor Sulfo-N-succinimidyl Oleate (SSO; 25μM) reduced the signal by almost half (46 (mean) ± 6% (SEM), one-way ANOVA, *p =* 1.41 × 10⁻⁸). Treatment with the microtubule inhibitor colchicine (0.2μg/ml) lowered the pHrodo Red area/ object area to 34 ± 6% (*p =* 2.98 × 10⁻¹²). Ivermectin (Importin α/β complex inhibitor; 1μM) treatment reduced nuclear pHrodo Red area/ object (μm^2^/object, averaged to myelin-fed positive control) to 39 ± 6% (*p =* 6.63 × 10⁻¹¹) (**Fig. 1f**), similar to nuclear import inhibitor wheat germ agglutinin (WGA; 100μg/ml), which resulted in a reduced area/ object to 36 ± 4% (*p =* 8.81 × 10⁻¹¹). In contrast, the nuclear exportin inhibitor leptomycin B significantly increased area/ object to 187 ± 17% (*p =* 8.0 × 10⁻¹⁵). The findings indicate that nuclear import of myelin proteins is phagocytosis-, microtubule- and importin-dependent.

### Five myelin mRNAs are enriched in the myelin-fed MG nuclei

To confirm translocation of myelin-derived mRNAs within MG, we used bulk RNA sequencing, comparing the profiles of myelin as well as post-fed MG fractions from cytoplasm and nucleus to unfed MG cytoplasm and nucleus fractions (**Fig. 2a**). We took advantage of species-specific differences to distinguish imported mouse versus rat myelin mRNAs. As shown (**Fig. 2b**, Extended Data **Tab 1**), of 19348 myelin mRNA species identified in myelin, five were enriched (>2.1-fold, DESeq2 Wald test, edgeR QLF test, limma-trend) within nuclei of myelin-fed MG. In rank order, these were: (1) *Mobp* (enriched 19.7-fold), (2) *Plekhb1* (enriched 4.9-fold) and (3) *Mbp* (enriched 3.5-fold) (**Fig. 2c**); (4) *Plp1* (2.5-fold) and (5) *Nfasc* (enriched 2.1-fold). We noted that *Mbp* and *Mobp* mRNAs were relatively abundant both in myelin and MG nuclei. In contrast, while *Plp1* was the third most abundant mRNA in the mouse myelin fed to MG (9.3 Log_2_CPM), it was less enriched (3.4 Log_2_CPM, 2.5-fold enriched in fed MG-nuc) relative to *Mobp* mRNA in the MG nuclei. These findings suggest that myelin RNAs are preferentially imported to nucleus versus cytoplasmic cell compartments. To confirm these findings, we conducted qPCR and smFISH on the myelin-fed MG nuclei. This showed that *Mobp* was enriched 7.0 (Log_2_FC) ± 1.8 (SD) fold (Welch ANOVA, *p =* 0.0174) (**Fig. 2d**), compared to unfed control, whereas *Mbp* in myelin-fed MG nucleus was enriched 2.3 ± 1.8 fold (*p =* 0.5593). We confirmed the presence of *Mbp* and *Mobp* transcripts within MG nuclei using sm-FISH (Extended Data **Fig. 3a, b**, 34% nuclei were *Mbp*+, compared to 4% of control, Welch’s t-test,; 16% of nuclei were *Mobp*+, compared to 3% of background, *p =* 1.0 × 10^⁻4^). These findings suggest the possibility of selective import of certain myelin mRNAs (e.g., *Mobp)* into the MG nucleus relative to others (e.g., *Plp)*. However, because *Mbp* and *Mobp* are also the most abundant species in myelin, non-selective processes are likely also involved.

**Figure 2.**
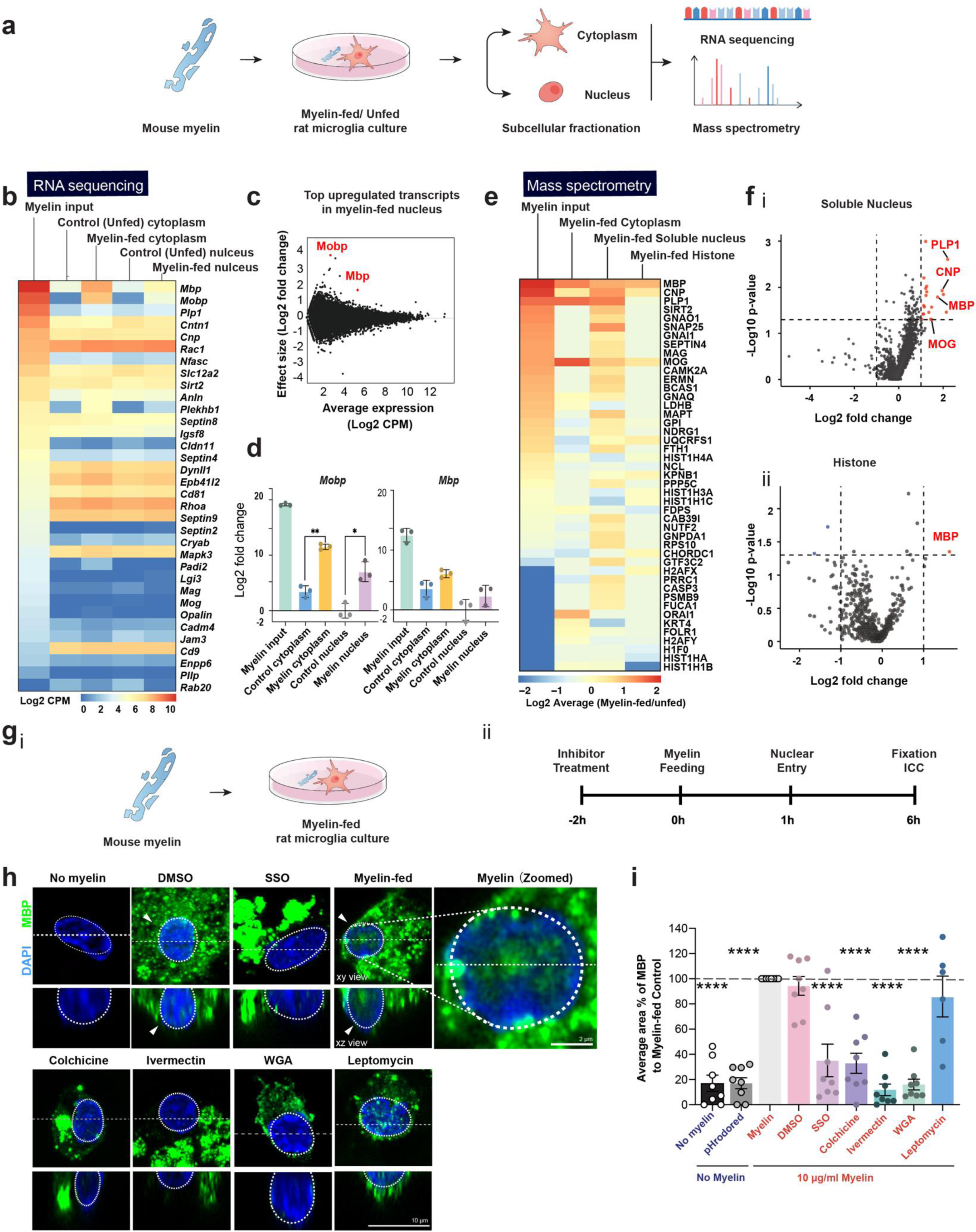
Omics analysis of myelin mRNA and protein import into MG nucleus. **a,** Schematic of the workflow for subcellular transcriptomic and proteomic analysis of myelin-fed microglia. Freshly prepared mouse myelin is fed to primary rat microglial cultures. At 24 h, cytoplasmic and nuclear fractions are isolated by subcellular fractionation and analysed by RNA sequencing and mass spectrometry. **b**, Heatmap of RNA-sequencing data showing enrichment of myelin-associated transcripts in the nuclear fraction of myelin-fed microglia. Data are presented as log₂ counts per million (CPM) for selected transcripts (*n =* 3). Columns (left to right): myelin input; cytoplasmic fractions (unfed and myelin-fed); nuclear fractions (unfed and myelin-fed). Myelin-related genes, including *Mobp* and *Mbp*, are enriched in the nuclear fraction following myelin feeding. Differential expression analysis was performed using the DESeq2 Wald test. The number of detected genes was 15,648 and 16,171 in nuclear RNA from control and myelin-fed microglia, respectively, and 15,747 and 16,106 in cytoplasmic RNA. c, MA plot of nuclear transcripts in microglia following myelin feeding. Effect size (log₂ fold change) versus average expression (log₂ CPM). *Mobp* and *Mbp* (red) are among the most strongly enriched transcripts in the nuclear fraction. **d**, RT–qPCR validation of nuclear enrichment of myelin-associated transcripts in microglia. Data are presented as mean ± s.d. (*n =* 3 independent biological replicates) of log₂ fold change relative to the unfed control. Relative expression of *Mobp* (left) and *Mbp* (right) is shown for myelin input, cytoplasmic and nuclear fractions from unfed and myelin-fed microglia. Statistical significance was assessed using Welch’s ANOVA followed by Games–Howell test for multiple comparisons versus the control. \**p <* 0.05, \*\**p <* 0.01 versus unfed nuclear control. **e,** Heatmap of proteomic data obtained by mass spectrometry showing log₂ fold change in protein abundance in myelin-fed versus unfed conditions across cytoplasmic and soluble nuclear fractions. Protein abundance in myelin (left column) is shown as log₁₀ average peptide counts. Myelin proteins, including MBP, PLP1, CNP and MOG, are enriched in the soluble nuclear fraction. MBP is the top enriched protein in the histone fraction. **f**, Volcano plots of differential protein enrichment in (i) the soluble nuclear proteome and (ii) the histone proteome of myelin-fed microglia relative to unfed controls. Data are presented as –log₁₀ P value versus log₂ fold change. Myelin-associated proteins (PLP1, CNP, MBP and MOG) are highlighted in the soluble nuclear fraction. MBP shows the greatest enrichment among myelin-associated proteins in the histone proteome. **g**, Schematic of imaging-based analysis of MBP nuclear entry. **i**, Myelin feeding in rat microglial cultures. **ii**, Timeline showing inhibitor pre-treatment (2 h before myelin feeding) and fixation for immunocytochemistry. **h**, High-magnification representative confocal immunofluorescence images showing nuclear localization of MBP following myelin feeding and fixation at 6 h. Single optical sections are shown; the lower panels show XZ sections along the dotted line. MBP (myelin basic protein; green) is detected within the nuclei of myelin-fed microglial cells (DAPI, blue; outlined), consistent with accumulation in the nuclear compartment and reduced by pharmacological inhibition (SSO, colchicine, ivermectin, wheat germ agglutinin and leptomycin B). Scale bars, 10 μm (left panels) and 2 μm (zoomed panels). **i**, Quantification of nuclear MBP accumulation (**h, ii**). Nuclear MBP-positive area is expressed as a percentage of the myelin-fed control and normalized to MBP⁺ microglial objects (MBP⁺ IBA1⁺ DAPI⁺ area, µm², per IBA1⁺ microglia). Data are presented as mean ± s.e.m. (*n =* 8). Each data point corresponds to one biological replicate, calculated as the normalized value averaged across >600 cells. Statistical analysis was performed using one-way ANOVA followed by Dunnett’s multiple comparisons test. Normality and homogeneity of variance were assessed using Shapiro–Wilk and Brown–Forsythe tests, respectively. \*\*\*\**p <* 0.0001 versus myelin-fed control. Nuclear MBP accumulation is reduced by pharmacological inhibition of CD36 (SSO), microtubules (colchicine), importin α/β (ivermectin) and nucleoporins (wheat germ agglutinin).

### MBP is singularly enriched in myelin-fed MG nuclear soluble and histone-bound fractions

We conducted quantitative tandem mass tag-mass spectrometry (TMT-M/S) to confirm translocation of myelin proteins within MG (**Fig. 2e)**. Myelin-fed MG were fractionated into cytoplasm, nuclear soluble fractions (**Fig. 2fi**) and nuclear histone (chromatin binding) fractions (**Fig. 2fii**). Amongst the top 15-enriched protein candidates in the nuclear soluble fraction, four known myelin structural proteins were significantly enriched (average FC difference > 1.5-fold, *p<* 0.05, -Log_10_ P value > 1.3), including PLP (4.6-fold, *p =* 0.00249), CNP (4-fold, *p =* 0.0118), MBP (3.2-fold, *p =* 0.00249) and MOG (2.8-fold, *p =* 0.0496). Another four myelin-associated/ regulatory proteins were also enriched: SIRT2 (2.6-fold, *p =* 0.0269), ERMN (2.6-fold, *p =* 0.0347), BCAS1 (2.5-fold, *p =* 0.0125) and FTH1 (2.3-fold, *p =* 0.0105) (Extended Data **Tab. 2**). Interestingly, this analysis also showed a significantly enriched endogenous KPNB1 (Importin subunit beta-1) (1.6-fold, *p =*0.0494) in the histone fraction, part of the nuclear pore complex (NPC). Of the 15 top-enriched proteins in the nuclear soluble fraction, only MBP was also enriched (3-fold, *p =* 0.0447) in the histone (chromatin-binding) nuclear fractions (**Fig. 2e,fii**). We validated MBP localization by antibody staining (**Fig. 2g, h**) in rat MG fixed at 6-hrs post-feeding (1µg/ml, 0.1ng/ cell). MG fed with myelin and treated with phagocytosis, microtubule and nuclear import inhibitors showed reduced MBP transport (**Fig. 2h, i**). These findings indicate that MBP was the sole myelin protein enriched in both soluble and histone-bound fractions of the MG nucleus. Thus, as the primary myelin protein translocated to the MG nucleus, it was selected for further evaluation.

### Purified myelin basic proteins rapidly access the MG nuclei

We next investigated if MBP is capable of cell and nuclear entry independent of myelin (**Fig. 3ai, ii**). As shown (**Fig. 3**), exogenous pHrodo Red labelled native pig MBP was rapidly detected in the nucleus 10 mins post-feeding (**Fig. 3b, Suppl. Video 3**), and signal continued to increase over 24 hrs. Analysis of human recombinant MBP yielded similar findings (Extended Data **Fig. 4**). The increase in nuclear signal intensity fed with pig or human MBP (from 1 to 24 hrs post-feeding) were respectively 4.0-fold (two-way ANOVA, *p =* 2.07 × 10⁻⁷) and 5.2-fold (*p =* 4.25 × 10⁻¹⁰) (**Fig. 3c**), higher than mouse myelin (2.4-fold, *p =* 4.45 × 10⁻⁶) or human myelin (2.6-fold, *p =* 6.22 × 10⁻⁵) and clear nuclear localization were observed (**Fig. 3e ii**, rendered with Imaris). We then investigated if nuclear entry of native pig MBP was blocked by inhibitors described above (**Fig. 3di, ii**). Similar to results obtained with untagged MBP (**Fig. 3e i**, Extended Data **Fig. 4**), SSO, colchicine, WGA and ivermectin all inhibited nuclear uptake (**Fig. 3e iii, 3f**). Interestingly, M9M, a specific peptide inhibitor of importin β2 subunit, greatly reduced MBP uptake (5.7 ± 1.7%, mean ± SEM) (**Fig. 3f**, one-way ANOVA, *p =* 1.35 × 10⁻¹²), suggesting an importin β2 subunit (transportin-1/ TNPO1)-dependent mechanism. To rule out that such effects were related to MBP toxicity, proteins were also assayed for endotoxin properties and MG cell death (Extended Data **Fig. 5**). These findings indicated that MBP accesses the MG nucleus via phagocytosis, intracellular and nuclear transport, independent of other myelin components.

**Figure 3.**
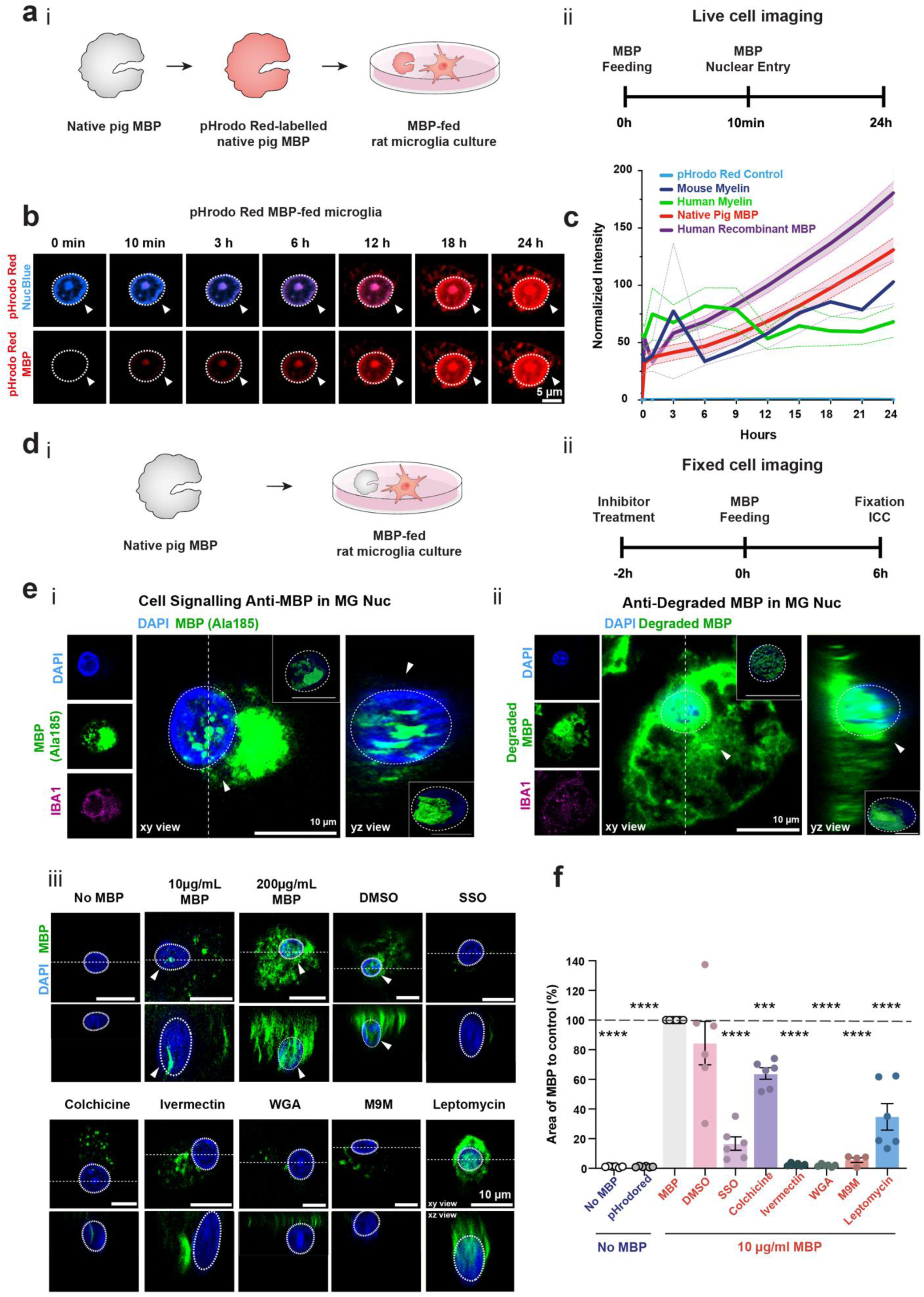
Myelin basic protein is retro-transported to MG nucleus *in vitro* by Importin α/β. **a,i**, Schemati of MBP feeding assays using pHrodo Red–labelled native pig MBP in microglia. i, pHrodo Red labelling of native pig MBP and feeding to CD11b⁺ rat microglia. **ii**, Timeline of live-cell imaging experiments to assess nuclear entry. b, Representative confocal time-lapse images of microglial cells following incubation with pHrodo Red–labelled MBP. Fluorescence is detected in the nucleus (outlined; NucBlue Live Cell nuclear stain) within 10 min of MBP addition and increases over 24 h. Scale bar, 5 μm. **c**, Quantification of normalized nuclear pHrodo Red intensity over time following incubation with mouse myelin, human myelin, native pig MBP or recombinant human MBP (mean ± s.e.m). Each data point represents the mean value per movie. Native pig MBP (red), human recombinant MBP (magenta). Statistical analysis was performed using two-way ANOVA followed by Dunnett’s multiple comparisons test. \*\*\*\**p <* 0.0001, for MBP and myelin-fed versus pHrodo red-fed control. **d,** i, Schematic of native pig MBP feeding in CD11b⁺ rat microglia. ii, Inhibitor pre-treatment (2 h before MBP feeding) followed by fixation at 6 h for analysis of nuclear MBP entry by immunocytochemistry. **e**, Representative confocal images showing nuclear MBP staining (green), detected using antibodies against **i**, MBP (Cell Signaling Technology (D8X4Q, #78896)) and **ii**, degraded MBP (anti-degraded MBP (Merck, AB5864)). Cells are fixed at 6 h following MBP feeding (200 μg ml⁻¹). Single optical sections are shown. Individual channels are displayed to the left for each condition, and orthogonal views are shown to the right along the dotted line. IBA1 staining identifies microglia. Imaris-generated 3D renderings corresponding to the displayed images are shown in the subpanel. Scale bar, 10 µm. **iii**, Representative confocal images showing nuclear MBP following treatment with 10 µg mL⁻¹ (83.33 pg per microglia) or 200 µg mL⁻¹ (1.6 ng per microglia) MBP. Nuclear accumulation of MBP is reduced upon treatment with the indicated intracellular transport inhibitors. Drug treatments were performed as in Fig. 2d,e. Inhibition of CD36-dependent uptake, microtubule transport, importin α/β1, importin β2, nucleoporin-mediated transport, or nuclear export reduces MBP nuclear localization. Scale bar, 10 µm. **f**, Quantification of inhibitor analysis shown in eii. Nuclear MBP-positive area is expressed as a percentage of the MBP-fed control and normalized to MBP⁺ microglial objects (MBP⁺ IBA1⁺ DAPI⁺ area, µm², per IBA1⁺ microglia). Data are shown as mean ± s.e.m. (*n* = 6). Each data point corresponds to one biological replicate, calculated as the normalized value averaged across >600 cells. Statistical analysis was performed using one-way ANOVA followed by Dunnett’s multiple comparisons test. Normality was assessed using the Shapiro–Wilk test, and homogeneity of variance using the Brown–Forsythe test. Adjusted P values are indicated *(***p <* 0.001). Colchicine (0.2 µg mL⁻¹; microtubule inhibitor) reduced nuclear MBP to 63.86 ± 9.19%. SSO (25 µ M; CD36-mediated phagocytosis inhibitor) reduced it to 16.64 ± 10.79%. Ivermectin (1 µ M; importin α/β inhibitor) reduced nuclear MBP to 2.57 ± 0.92%. Wheat germ agglutinin (100 µg mL⁻¹; nucleoporin/glycoprotein inhibitor) reduced it to 1.90 ± 0.89%. M9M (10 µ M; importin β2 inhibitor) reduced nuclear MBP to 5.73 ± 1.71%.

### MBP is an RNA-binding protein

Although the MBP 21.5-kDa isoform contains a non-traditional PY-type nuclear localization signal (**Fig. 4a**) ^39,45,46^, which accounts for importin-dependent transport, the RNA binding functions of MBP have not been described. To assess this, we first performed a dot blot assay using native pig MBP and tagged (biotinylated) *Mobp* mRNA (0–20 nM). MBP showed a concentration-dependent retention on the membrane, indicating *Mobp* mRNA binding (**Fig. 4b**). The dissociation constant (Kd) was calculated against a temperature gradient using the Hill–Langmuir equation, yielding a value of 0.2967 nM (Hill coefficient = 2; T = 45.5°C; T = 60°C; L =2 nM). The melt curve analysis thus indicated a high-affinity binding between MBP and *Mobp* mRNA (**Fig. 4c**).

**Figure 4.**
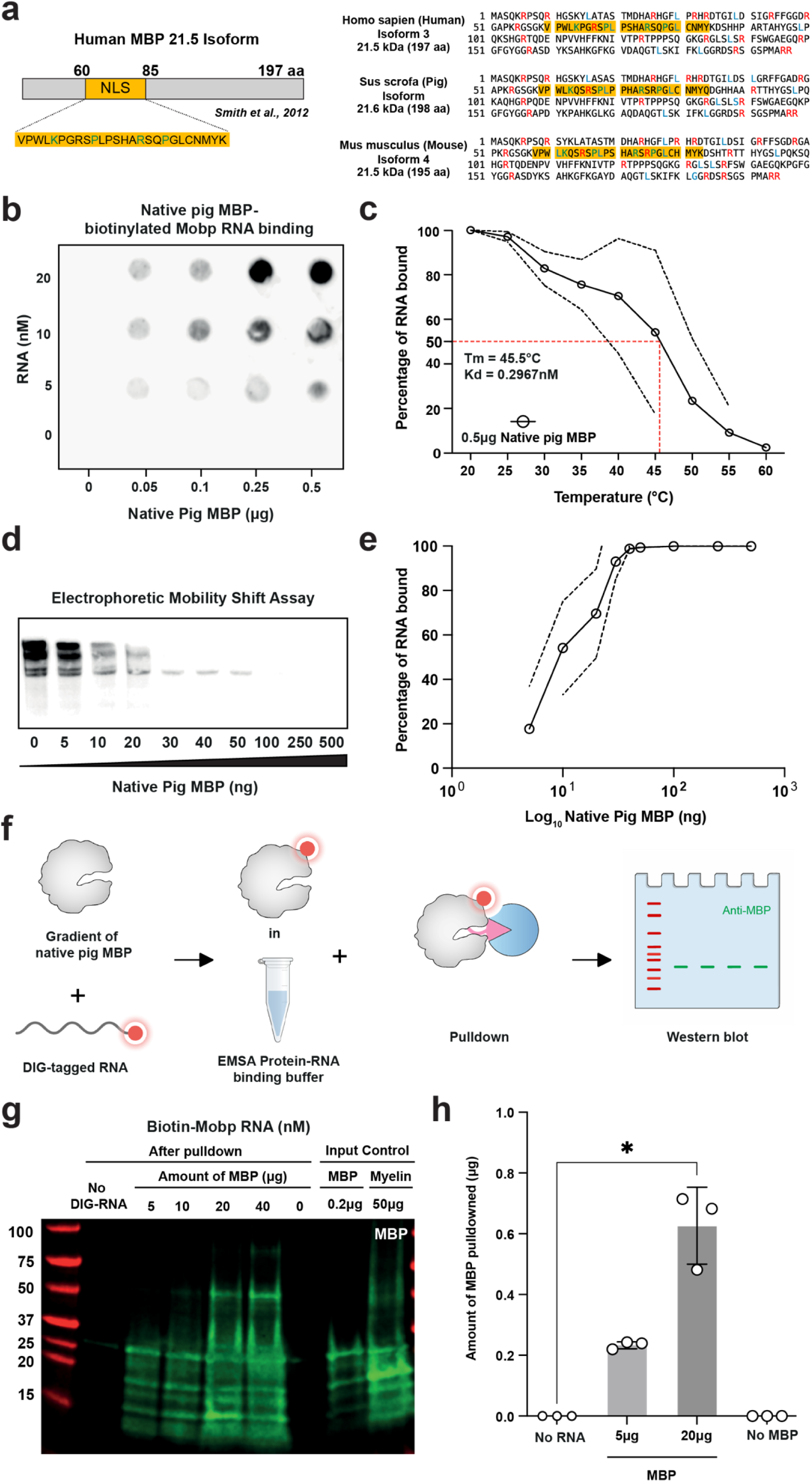
MBP is a RNA-binding protein. **a**, Schematic of the human MBP 21.5-kDa isoform and conservation of the N-terminal NLS/RNA-binding region. Left, schematic representation of the human MBP 21.5-kDa isoform, highlighting the nuclear localization signal (NLS; yellow) proximal to the N-terminus. Right, sequence alignment of MBP from human, pig, and mouse, showing high conservation of basic residues within the NLS/RNA-binding region. **b,** Dot-blot binding assay showing RNA binding to immobilized native pig MBP. Increasing concentrations of RNA (5–20 nM) were incubated with MBP spotted at the indicated amounts (0–0.5 µg), and binding was detected using biotinylated *Mobp* RNA. **c,** Thermal denaturation analysis of MBP–RNA binding (n = 3). The fraction of RNA bound (%) is plotted as a function of temperature, showing progressive loss of binding with increasing temperature. Curve fitting yields an apparent melting temperature (Tₘ) of ∼45.5 °C and an estimated dissociation constant K_d_) of ∼0.30 nM, consistent with high-affinity interaction. **d**, Electrophoretic mobility shift assay (EMSA) showing dose-dependent binding of native pig MBP to 20 nM biotin-labelled *Mobp* RNA. Increasing MBP concentrations result in a progressive shift of RNA. At 100ng MBP, RNA is fully shifted, indicating complete binding. The unbound fraction of biotinylated RNA decreases with increasing MBP, as shown by the diminishing free RNA band. **e,** Quantification of EMSA data showing saturation of RNA binding with increasing MBP concentrations (*n* = 3). The fraction of 20 nM biotin-labelled *Mobp* RNA bound is plotted as a function of MBP amount. **f,** Schematic of the DIG–RNA–protein pulldown workflow. DIG-tagged RNA was incubated with increasing amounts of native pig MBP, followed by pulldown and analysis by western blot. **g**, Western blot analysis of MBP recovered following DIG–RNA pulldown. MBP enrichment increases with DIG-labelled RNA and increasing MBP input, compared with no-RNA controls. Input controls are shown on the right. Lanes: 1, molecular weight marker; 2, no DIG–RNA + 10 µg MBP; 3, 5 µg MBP; 4, 10 µg MBP; 5, 20 µg MBP; 6, 40 µg MBP; 7, RNA only (no MBP); 8, input control (0.2 µg MBP); 9, input control (50 µg Myelin). **h**, Quantification of MBP pulldown efficiency (n = 3). MBP recovery is increased in RNA-containing pulldowns relative to no-RNA controls. Data are shown as mean ± s.d. Statistical analysis was performed using the Kruskal–Wallis test followed by Dunn’s multiple comparisons test (*P < 0.05).

We next used gradient limitation assays in the form of electrophoretic mobility shift assays (EMSAs) to evaluate MBP’s RNA binding capacity, whereby unbound (biotinylated) mRNA show as bands in the blot. As shown (**Fig. 4d, e**), *Mobp* mRNA-MBP complexes displayed reduced mobility compared to unbound RNA. High MBP concentrations (0.5–0.1 μg) bound most of the *Mobp* mRNA signal (and thus were absent on the gel), whereas at lower MBP concentrations (0.05–0.005 μg), progressively more unbound *Mobp* mRNA was detected. RNA pull-down assays further confirmed physical interaction between MBP and *Mobp* mRNA (**Fig. 4f**). Digoxigenin (DIG)-tagged *Mobp* mRNA (-UTP-labelled) incubated with MBP followed by anti-DIG bead enrichment resulted in concentration-dependent co-purification of MBP, plateauing at 20 μg MBP (**Fig. 4g–h**) (Kruskal-Wallis test, *p =* 0.0196). Together, these findings indicate MBP is a nucleoprotein capable of binding mRNA.

### MBP functions as an RNA chaperone for nuclear retro-transport

We next evaluated whether MBP was sufficient for RNA retro-transport into the nucleus of MG *in vitro*. Using DIG-tagged *Mobp* mRNA and either native pig or recombinant human MBP (**Fig. 5a–b**), we detected puncta of DIG-mRNA within MG nuclei in 6 hrs post-feeding (one-way ANOVA, *p =* 1.41 × 10⁻¹¹) (**Fig. 5bi, iii, iv**). Treatment with import inhibitors significantly decreased nuclear RNA localization (**Fig. 5bii**). Specifically (**Fig. 5c**), ivermectin (1 μM; importin α/β inhibitor) reduced nuclear RNA to 18 ± 4% (SEM) (*p =* 1.91 × 10⁻⁹), wheat germ agglutinin (100 μg/mL; functions to obstruct the nuclear pore transport channel, also inhibits Importin α/β–mediated import) reduced it to 42 ± 7% (*p =* 5.09 × 10⁻⁵), and the microtubule inhibitor colchicine (0.2 μg/mL) and CD36 inhibitor SSO (25 μM) caused moderate decreases of 63 ± 8% (*p =* 0.0198) and 56 ± 8% (*p =* 0.00396). These results suggest that both microtubules and the nuclear pore complex contribute to MBP-mediated RNA transport.

**Figure 5.**
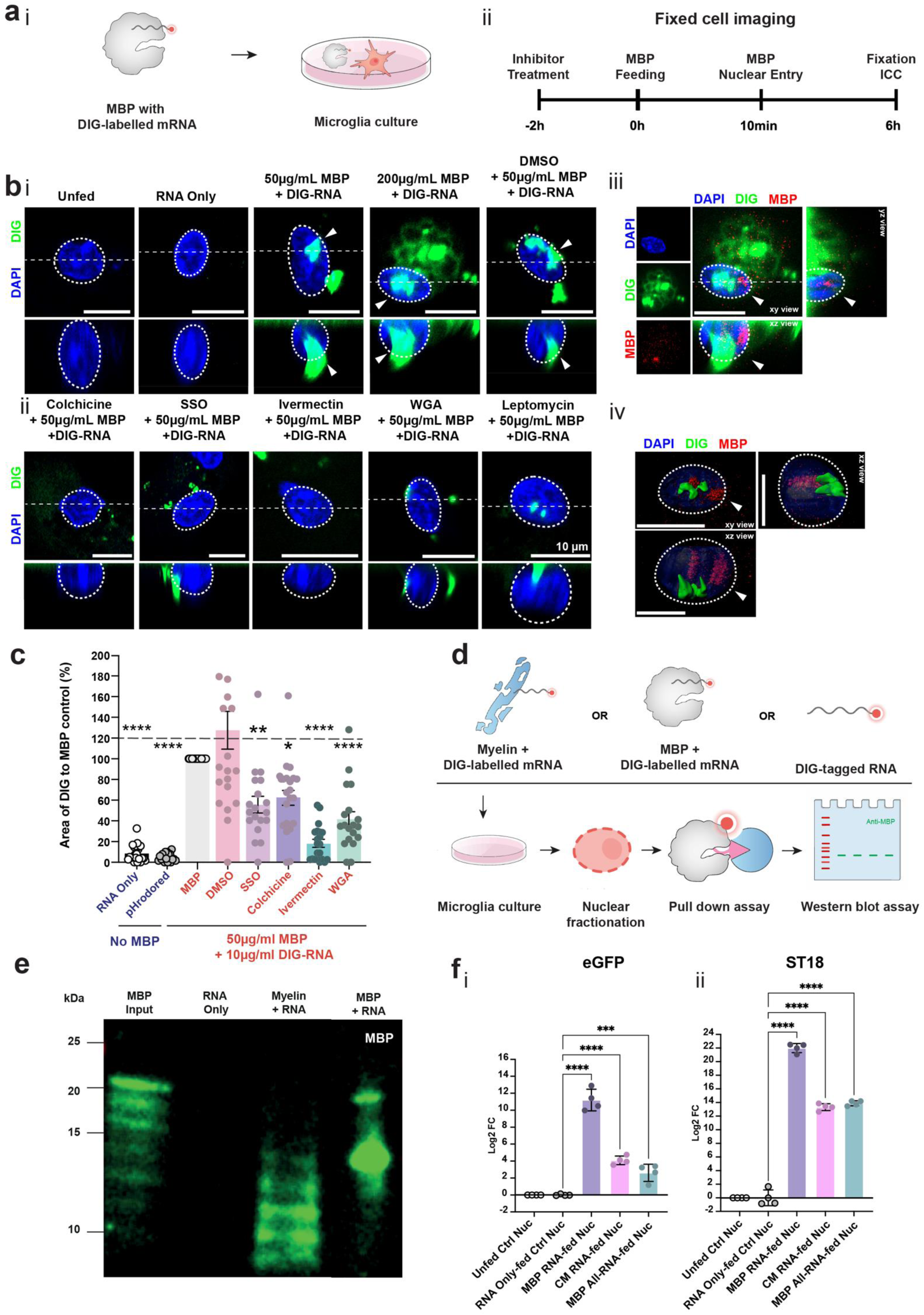
MBP is a RNA chaperone for non-specific RNA retro-transport to the MG nucleus *in vitro*. **a,i,** Schematic illustrating MBP-dependent nuclear delivery of RNA in primary CD11b⁺ rat microglial cultures. mRNA was pre-labelled with digoxigenin (DIG-UTP). **a**,**ii**, Timeline of fixed-cell imaging experiments to assess nuclear entry. Inhibitors were applied 2 h before myelin feeding, and cells were fixed at 24 h for ICC to assess nuclear RNA entry. **b**,**i**, Representative confocal images of nuclear RNA delivery in microglia. Nuclear-delivered DIG-labelled RNA (green; arrowheads) is detected within nuclei (DAPI, blue; outlined by dashed white lines). Single optical sections are shown. The lower panel shows an orthogonal XZ view along the dotted line. Scale bar, 10 μm. Conditions (left to right): unfed, RNA only, 50 μg ml⁻¹ MBP + DIG-labelled RNA, 200 μg ml⁻¹ MBP + DIG-labelled RNA, and vehicle (DMSO) control. b,ii, Representative confocal images illustrating the effects of pharmacological inhibitors (colchicine, SSO, ivermectin, wheat germ agglutinin (WGA) and leptomycin B) on MBP-mediated nuclear accumulation of DIG-labelled RNA. Inhibitors were applied as described in Fig. 2d,e. Scale bar, 10 µm. **b,iii**, Representative confocal images of nuclear RNA delivery in microglia exposed to 200 µg ml⁻¹ MBP together with DIG-labelled RNA. Nuclear-localized DIG-labelled RNA (green; arrowheads) is detected within DAPI-stained nuclei (blue; outlined by dashed white lines), with MBP shown in red. Single optical sections are shown. Lower panels display orthogonal XZ views along the dotted line. Scale bar, 10 µm. **b**,**iv**, Imaris-generated three-dimensional reconstruction of nuclear-imported DIG-labelled RNA in microglia exposed to 200 µg ml⁻¹ MBP together with DIG-labelled RNA. DAPI (blue) marks nuclei, DIG-labelled RNA (green) highlights nuclear-localized RNA, and MBP is shown in red. Orthogonal views (xy, xz and yz) are shown. Scale bars, 10 µm. **c**, Quantification of nuclear accumulation of DIG-labelled RNA in MBP-fed and drug-treated conditions. Nuclear area positive for DIG signal is expressed as a percentage of the MBP-fed control. Data are presented as mean ± s.e.m. (*n =* 21). Each data point corresponds to one biological replicate, calculated as the normalized value averaged across >600 cells. Statistical analysis was performed using one-way ANOVA with Dunnett’s multiple comparisons test. Normality was assessed using the Shapiro–Wilk test, and equal variance was assessed using the Brown–Forsythe test. P-value summaries are indicated as follows: \**p <* 0.05; \*\**p <* 0.01; \*\*\**p <* 0.001;\*\*\*\**p <* 0.0001. **d**, Schematic of the anti-DIG pulldown and western blot workflow to assess MBP–RNA interactions *in vitro*. Microglia were exposed to myelin + DIG-labelled RNA, MBP + DIG-labelled RNA, or DIG-labelled RNA alone. This was followed by nuclear fractionation, anti-DIG pulldown, and analysis by western blot. **e**, Representative western blot showing MBP recovery following RNA pulldown. Samples include RNA alone, myelin + RNA, and MBP + RNA, compared with MBP input control. f, RT–qPCR analysis of exogenous RNA enrichment in microglial nuclei. **i**, Nuclear levels of exogenous *egfp* mRNA following RNA-only delivery, MBP–RNA co-delivery, myelin–RNA co-delivery, or MBP-alone treatment. Data are presented as log₂ fold change relative to the RNA-only nuclear control (mean ± s.d.; *n* = 4). Statistical analysis was performed using one-way ANOVA followed by Dunnett’s multiple comparisons test against the RNA-only control. Adjusted *p* values are indicated (\*\*\**p* < 0.001, \*\*\*\**p* < 0.001). **ii**, Nuclear levels of exogenous *ST18* mRNA under the same conditions.

To validate RNA–protein complex formation during delivery into the nucleus, we fractionated the nuclei and used anti-DIG (beads) pulldown and western blots to further validate the RNA-protein binding affinity (**Fig. 5d, e**). Gel electrophoresis of the pulled down proteome indicated that when myelin or MBP were co-fed with DIG-tagged RNA, MBP was recovered with anti-DIG beads (**Fig. 5e**). We also found that untagged *Mobp* mRNA was also trafficked into MG similarly (Extended Data **Fig. 6a, b)**; interestingly, we found the RNA cargo also remained intact and could be amplified with PCR (Extended Data **Fig. 6ci, ii**). These findings indicate that MBP is an RNA-binding/trafficking protein for MG nuclear import *in vitro* and suggests it also preserves mRNA cargo for at least 48 hours post-ingestion.

While our data provided evidence for preferential RNA trafficking to the nucleus (**Fig. 2b, 5b**), it was possible that MBP bound RNA non-selectively (e.g., through electrostatic interactions). Using orthologous *egfp* mRNA (1.32 kb) and oligodendrocyte mRNA transcript *st18* (3.6 kb), we found both (*egfp* (Log_2_FC =11 ±1, one-way ANOVA, *p =* 7.24 × 10⁻¹²); *st18* (Log_2_FC =22 ± 0.7, *p<* 1 × 10⁻¹⁵) were significantly enriched in MG nuclei when co-incubated with MBP (**Fig. 5fi,ii**). Competitive binding assays with co-administered *egfp, st18* or random RNAs revealed reduced enrichment of both *egfp* (Log_2_FC =2.6 ± 1) (*p =* 0.000785) and *st18* (Log_2_FC =14 ± 0.5) (*p =* 3.7 × 10⁻¹⁴) transcripts, suggesting substrate competition for MBP binding (**Fig. 5f**). These findings suggest that MBP can bind a broad range of mRNAs through low-specificity protein-nucleic acid interactions.

### MBP acts as a chaperone for mRNA retro-transport into MG nuclei *in vivo*

To further validate our findings, we asked whether MBP would target MG and deliver RNA to the MG nuclei *in vivo*. We first stereotaxically injected pHrodo Red-labelled MBP into the brains of mice at age P8. 5 days post-injection (d.p.i.) (*n =* 6), nuclear pHrodo Red signal colocalized with dMBP immunolabeling (Millipore, cat. #AB5864^93^, 1:300), using an antibody that detected pig-MBP from endogenous mouse MBP (**Fig. 6a-c**). As indicated (**Fig. 6d**), the vast majority (92% ± 11%) of nuclear pHrodo Red⁺ puncta co-localized with degraded MBP within Iba1⁺ cells. To test whether injected MBP facilitates mRNA retro-transport to MG nuclei, we injected DIG-*Mobp* mRNA with either 3μg of MBP or MAG (a transmembrane myelin protein localized to the periaxonal membrane ^94^) controls (*n =* 3, for each group). Using high-resolution confocal microscopy and deconvolution, we quantified retro-transported mRNA puncta in the brain cells (**Fig. 6e-g, Extended Data Fig. 7)**. We quantified the injection site and the surrounding area, dividing cells of the parenchyma into MG and non-microglia (non-MG) cells. This indicated that co-injection of MBP (but not MAG) with mRNA significantly increased DIG particles in MG nuclei (**Fig. 6g, 6hi,, Extended Data Fig. 7**) (1.5 ± 0.4 DIG+ particles in mRNA only vs 1.5 ± 0.3 DIG+ particles in mRNA+MAG vs 11± 2 DIG+ particles in mRNA+MBP, nested ANOVA with multiple comparisons, *p value* = 0.0075). By comparison, non-MG showed no significant difference between conditions (**Fig. 6g, 6hii,, Extended Data Fig. 7**) (0.4 ± 0.04 DIG+ particles in mRNA only vs 2.2 ± 0.2 DIG+ particles in mRNA+MAG vs 3.7 ± 0.4 DIG+ particles in mRNA+MBP). Furthermore, we quantified the proportion of cells in each population containing 10 or more DIG+ particles. We found that within non-MG cells, there was no significant increase in the presence of MBP (0.2 ± 0.2% positive in mRNA only vs 4 ± 2.2% positive in mRNA+MAG vs 11 ± 1.3% positive in mRNA+MBP), while a significant proportion of MG was positive for DIG+ mRNA (2 ± 2% positive in mRNA only vs none positive in mRNA+MAG vs 36 ± 12% positive in mRNA+MBP, 2way ANOVA, *p =* 1.13 × 10⁻⁵ (**Fig. 6i**). Together, these data show that, *in vivo*, MBP can act as a chaperone for MG nuclear retro-transport of mRNAs. A limitation of IBA1⁺ cell identification in this study is that this marker is not exclusive to resident microglia and may also label infiltrating macrophages, thereby precluding clear discrimination between these populations.

**Figure 6.**
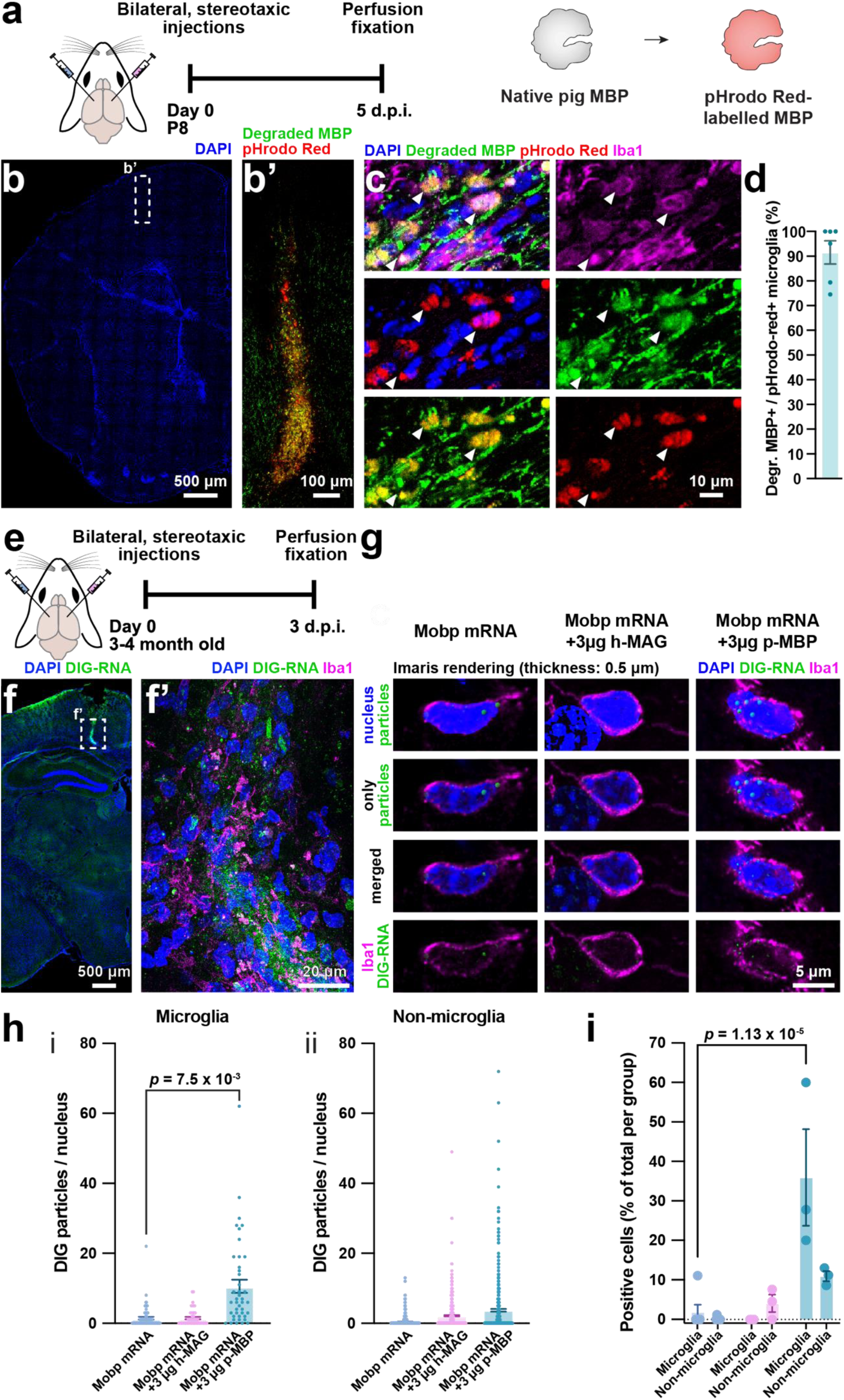
MBP is a RNA chaperone for RNA retro-transport to the MG nucleus *in vivo*. **a.** Schematic overview of the *in vivo* nuclear MBP uptake assay in P8 mouse pups. **b**. Representative brain hemi-section of an injected animal. Nuclei labelled with DAPI (blue). Box indicates the injection site. **b’** Magnified view of the injection site, labelled with degraded MBP (green) and pHrodo Red (red). Scale bars: 500 and 100 μm respectively. **c**. High-magnification micrograph of the injection site indicating nuclei (DAPI, blue) labelled with pHrodo Red (red) and degraded MBP (green). Microglia were identified by Iba1⁺ staining (magenta). Example nuclei are outlined. Scale bar: 10 μm. **d**. Quantification of degraded MBP⁺ population within pHrodo Red+ve microglia cells. Data shown as mean±SEM (*n =* 6). **e.** Schematic overview of the *in vivo* nuclear MBP uptake assay in 3-4 month-old mice. **f.** Representative brain hemi-section of an injected animal. f’, Magnified view of the injection site, labelled with DIG+ve RNA (green), Iba1 (magenta) and DAPI (blue). Scale bars: 500 and 20 μm respectively. **g.** Representative 0.5 μm thick Imaris-rendered micrographs of microglia cell bodies (Iba, magenta) from *Mobp* mRNA (left), *Mobp* mRNA + 3 µg h-MAG (middle) or *Mobp* mRNA + 3 µg p-MBP (right) injected animals. Segmented nuclei are rendered in blue (DAPI), while DIG+ve particles within nuclei are rendered as spheres (green). Scale bar: 5 μm. **h**. Quantification of DIG+ve particles in the nucleus in (**i**) microglia (left) and in (**ii**) non-microglia cells (right) across conditions. Data are shown as mean±SEM (*n =* 3). Increased, selective uptake was observed in Iba1+ve cells with 3 µg p-MBP + *Mobp* mRNA. Statistical analysis was performed using nested one-way ANOVA with Dunnett’s multiple comparisons test (vs *Mobp* mRNA control). Normality was assessed using the Shapiro–Wilk test, and equal variance was assessed using the Brown–Forsythe test. **i**. Quantification of the proportion of cells that are DIG+ve over the total number of cells in their respective type (microglia vs non-microglia). Data are shown as mean±SEM. Increased proportion was seen only in microglia and only in the presence of 3μg of p-MBP. 2-way ANOVA with Dunnett’s multiple comparisons test (vs *Mobp* mRNA control). ANOVA P value < 0.0001.

### MBP facilitates nucleic acid uptake to regulate MG inflammatory responses

The studies above addressed mechanisms of MBP mRNA transport into MG and potential roles in the nucleus in bound and soluble histone fractions. We next tested MG cellular regulation via knockdown of *TLR4*, a sensor in the innate immune response to pathogen-associated molecular patterns (PAMPs) and damage-associated molecular patterns (DAMPs) (**Fig. 7ai**), with functions in activation and secretion of pro-inflammatory cytokines TNF-α, IL-6, IL-1β and CXCL10 ^95^. As shown (**Fig. 7aii**), we achieved knockdown with 30nM siRNA delivered using MBP protein-to-siRNA mass ratios ranging from 10:1 to 100:1, with efficiencies of 30-45% compared to negative controls. Efficiencies were 30 ± 18% (10:1) (one-way ANOVA, *p =* 0.0117), 34 ± 17% (50:1) (*p =* 4.90 × 10⁻⁹), and 34 ± 22% (100:1) (*p =* 1.33 × 10⁻⁴). At a higher siRNA concentration of 50 nM, knockdown efficiencies yielded 49 ± 27% (*p =* 4.90 × 10⁻⁴). For comparison, lipofectamine-based controls resulted in knockdown efficiencies between 47 ± 19 % (30nM) (*p =* 1.43 × 10⁻⁴) and 44 ± 16% (50nM) (*p =* 2.02 × 10⁻⁵).

To assess the functional significance of TLR4 knockdown, we tested this against exposure to lipopolysaccharide (LPS) at a concentration of 5 ng/mL 72 hours post-knockdown for a timespan of 12 hours (**Fig. 7b**). Prior to LPS stimulation, we exposed MG to various treatment conditions, including MBP only, negative siRNA only, negative siRNA combined with MBP and siRNA at 30nM and 50nM combined with MBP.

**Figure 7.**
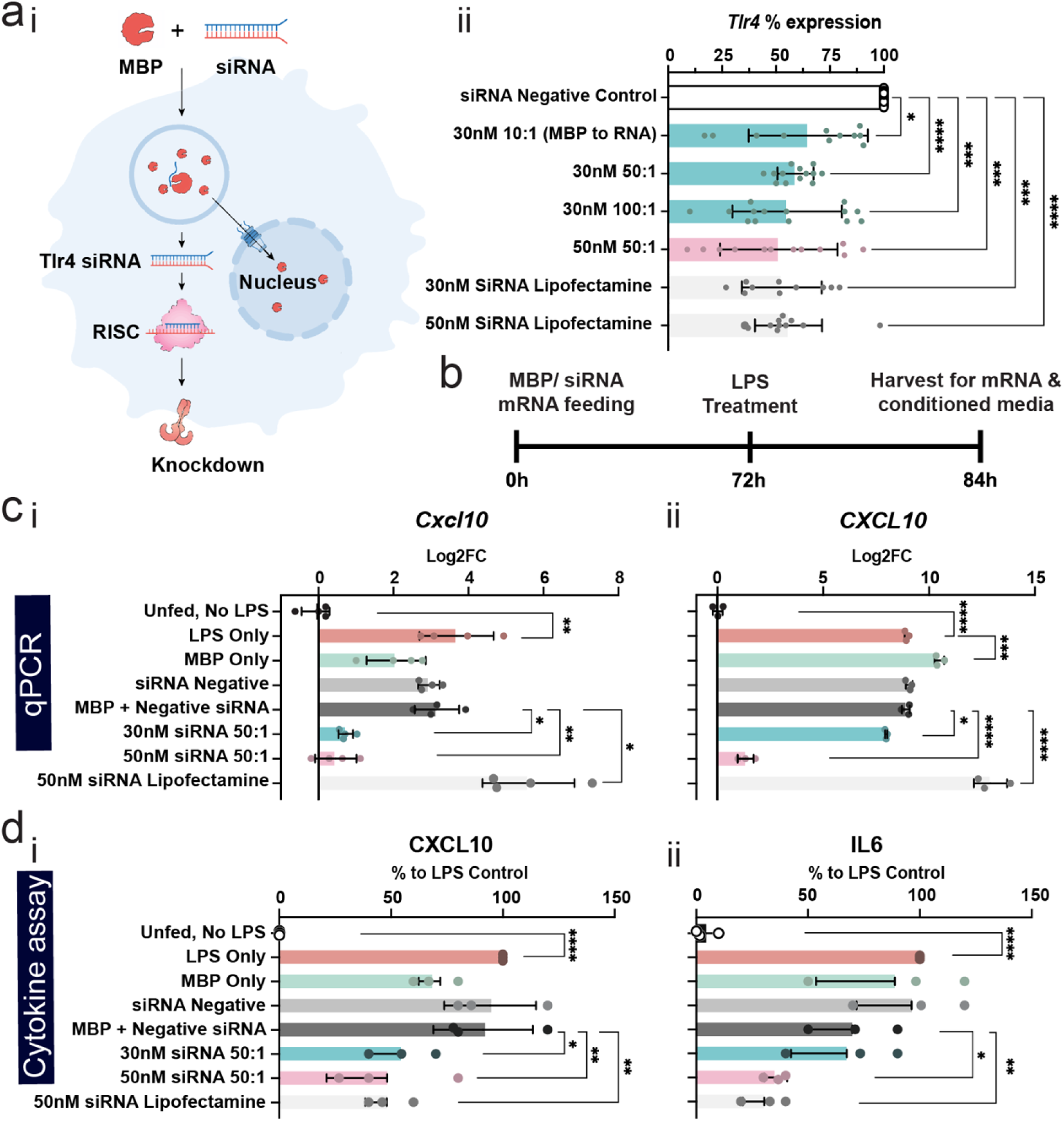
MBP attenuates MG inflammatory responses in an RNA-dependent manner by promoting siRNA-mediated TLR4 silencing. **a,i,** Schematic illustrating uptake of MBP–siRNA complexes by fed microglia, followed by RNA interference via RISC-mediated degradation of target *Tlr4* transcripts. **a,ii**, Quantification of *Tlr4* mRNA expression in primary mouse microglia following siRNA-mediated knockdown. Expression levels are shown as a percentage of the siRNA negative control. Cells were treated with MBP–siRNA complexes (30 nM or 50 nM) and compared with lipofectamine-positive controls. RNA was collected 36 h after treatment. Data are presented as mean ± s.d. (*n =* 15), with individual data points shown. Statistical significance was determined using one-way ANOVA with Dunnett’s multiple comparisons test (versus siRNA negative control). Normality of residuals was assessed using the Shapiro–Wilk test, and homogeneity of variances using the Brown–Forsythe test. Adjusted P values are indicated (\**p <* 0.05, \*\**p <* 0.01, \*\*\**p <* 0.001, \*\*\*\**p <* 0.0001). **b**, Timeline of MBP–Tlr4 siRNA-mediated knockdown in microglia at 0 h, stimulated with lipopolysaccharide (LPS) at 72 h and harvested at 84 h for mRNA quantification and conditioned media collection. **c**, RT–qPCR analysis of MBP-mediated Tlr4 knockdown attenuates LPS-induced inflammatory gene expression in (**i**) mouse and (**ii**) human microglia. LPS was applied to all conditions except the unfed, no-LPS control at 72 h post-knockdown. Log₂ fold change (log₂FC) in *Cxcl10* expression is shown for untreated, LPS-only, MBP-only, siRNA negative control, and MBP–siRNA-treated microglia, relative to the MBP + siRNA negative control. Data are presented as mean ± s.d. (*n =* 4 independent biological replicates), with individual data points shown. Statistical significance was determined using nested one-way ANOVA with Dunnett’s multiple comparisons test (versus MBP + siRNA negative control). Normality of residuals was assessed using the Shapiro–Wilk test, and homogeneity of variances using the Brown–Forsythe test. \**p <* 0.05, \*\**p <* 0.01, \*\*\**p <* 0.001, \*\*\*\**p <* 0.0001. **d**, MBP-mediated Tlr4 knockdown attenuates LPS-induced cytokine secretion in cultured mouse microglia. Cytokine levels in conditioned media were quantified for (**i**) CXCL10 (left), (**ii**) IL-6 (right). Experimental conditions are as described in panel f. Data are presented as mean ± s.d. (*n =* 3 independent biological replicates), with individual data points shown. Statistical significance was determined using ordinary one-way ANOVA with Dunnett’s multiple comparisons test (versus MBP + siRNA negative control). Normality of residuals was assessed using the Shapiro–Wilk test, and homogeneity of variances using the Brown–Forsythe test. \**p <* 0.05, \*\**p <* 0.01, \*\*\**p <* 0.001, \*\*\*\**p <* 0.0001

As shown (**Fig. 7ci**, *n =* 4), pretreatment with MBP-*Tlr4* siRNA at 30 nM and 50 nM significantly reduced *Cxcl10* mRNA expression (following LPS stimulation (5 ng/ml)) in mouse MG. 30 nM siRNA suppressed *Cxcl10* expression, yielding a reduction of Log2 2.3 ± 1.1 fold (nested ANOVA with multiple comparisons, *p =* 0.0484, normalized to MBP + negative siRNA control). 50 nM Tlr4 siRNA delivered with MBP resulted in a Log2 2.7 ± 1.3 fold reduction (*p =* 0.0082). Thus, in mouse MG, both 30 nM and 50 nM siRNA attenuates LPS-induced inflammatory Cxcl10 expression, via *Tlr4* knockdown. To validate fidelity across species, we also tested the knockdown in human MG (**Fig. 7cii**, *n =* 3). 30 nM siRNA delivered with MBP reduced *Cxcl10* expression by Log2 –0.98 ± 0.35 fold (nested ANOVA with multiple comparisons, *p =* 0.048, normalized to MBP + negative siRNA control). At 50 nM siRNA delivered with MBP, a pronounced Log2 –7.6 ± 0.6 fold reduction (*p =* 4.0 × 10⁻¹⁵) was resulted.

The attenuated inflammatory response was further validated by cytokine measurements. 50 nM siRNA delivered with MBP significantly reduced CXCL10 (Fig **7di**, *n =* 3) (0.44-fold to the MBP + siRNA negative control, one-way ANOVA, P = 0.001) and IL-6 expression (0.31-fold, P = 0.0309) (Fig **7dii**, *n =* 3). These findings indicate that efficient *Tlr4* knockdown is facilitated by MBP and that MBP can deliver RNA molecules that functionally impact microglial function.

## Discussion

In multiple sclerosis (MS), autoimmune-mediated oligodendrocyte damage generates abundant myelin debris that is phagocytosed by resident microglia (MG) and infiltrating macrophages, and several studies have shown myelin-derived mRNAs within MG nuclei ^60,61,67^. These observations raised the question of how ingested myelin-derived mRNAs were imported to the nucleus? We found that the prevalent myelin component MBP promotes RNA nuclear import in MG after phagocytosis of myelin debris. We show MBP is a nuclear chaperone capable of binding nucleic acids and mediating retrograde nuclear transport via importin-dependent mechanisms in MG. To our knowledge, this is the first report of a non-viral mechanism for retrograde nuclear mRNA transport.

### Retrograde transport of myelin components to MG Nuclei

Our *in vitro* experiments using fluorescently labelled myelin proteins confirmed rapid nuclear import of myelin macromolecules, a process that was inhibited by the importin inhibitors ivermectin ^84–86^ and wheat germ agglutinin (WGA) ^87,88^. Although both wheat germ agglutinin (WGA) and ivermectin exert non-specific inhibitory effects—WGA broadly disrupts nuclear pore–mediated transport, whereas ivermectin inhibits importin α/β–dependent nuclear import and indirectly affects NF-κB and interferon signalling—they remain useful tools for investigating nuclear import mechanisms. In this context, they were selected to demonstrate nuclear import of myelin proteins and associated mRNA because they provide robust inhibition of nuclear transport. Nuclear RNA sequencing of MG after myelin uptake revealed a distinct subset of myelin transcripts—including *Mbp*, *Mobp*, *Plp1*, *Nfasc*, and *Plekhb1*—comprising a small but reproducible fraction (0.056%) of total nuclear RNA. Parallel tandem mass tag proteomics of nuclear soluble fractions showed prominent nuclear enrichment of myelin proteins (MBP, PLP1, CNP, MOG), with MBP uniquely enriched in the nuclear chromatin fraction along with importin subunit beta-1, further supporting involvement of the nuclear transport machinery. The disproportionate transport of certain myelin mRNA and protein species warrants further mechanistic investigation.

### MBP properties, RNA-binding and nuclear trafficking

Originally sequenced in 1970 ^96,97^, myelin basic protein (MBP) is small (14-21.5KDa, isoforms-dependent)^39^, highly basic (Lys/Arg/His ∼25%) ^38^, strongly positively charged (pI∼10)^98^, of low hydrophobicity ^44^ and exhibits high intramolecular electrostatic repulsion ^42^. MBP has five isoforms in mice and four in humans ^39,45,46^, with the 18.5kDa and 21.5kDa isoforms conserved across both species ^35,39^. The 18.5kDa is most abundant in the human adult CNS (one of the major isoforms in mouse CNS)^99^, whereas the 21.5kDa isoform^45^ is more prevalent in oligodendrocytes development, myelin compaction and ^45,47^ and disease^48^.

The ssRNA influenza A virus^100,101^releases its viral ribonucleoproteins (vRNPs) into the cytoplasm that engage importin-α/β^77,78,101^, enabling active transport through nuclear pore complexes into the nucleus ^100,102,103^. Nuclear RNA import has also been observed for non-coding RNAs ^104^ and RNA–protein complexes^105^ (e.g., spliceosomal snRNPs^106^), telomerase RNA^107^ and distinct tRNAs^108^ under stress conditions ^109^. In these cases, RNA species re-enter the nucleus via importin-dependent mechanisms mediated by NLS-bearing chaperone proteins^79^. While prior work identified that the 21.5 kDa MBP isoform has a NLS-PY sequence^45^ that can promote nuclear uptake in HeLa^47^ and oligodendrocytes^46^, the biological significance of this has been unclear since the vast majority of MBP proteins are cytoplasmically located in myelin of oligodendrocytes. Here we show that 21.5-kDa MBP has a novel capability as an RNA-chaperone that efficiently shuttles myelin-associated mRNAs (*Mobp, st18*) or heterologous reporter transcripts (*egfp*) to the MG nucleus. We speculate that nonspecific mRNA binding is achieved through electrostatic interactions with positively charged amino acids, along with a putative glycine-phenylalanine-glycine (GFG)-RNA binding domain ^110^, although this remains to be tested. We found that MBP is phagocytosed *in vivo* and *in vitro* and delivers tagged RNA to the MG nucleus, and that terminal nuclear translocation was inhibited by importin α/β inhibitors ivermectin and WGA.

### Implications of MBP-RNA trafficking for neuroinflammation and gene engineering

The novel finding that MBP is a nucleoprotein chaperone for RNA provides a mechanistic explanation for how myelin mRNA transcripts become localized to the nuclei of phagocytes from human MS patients ^60,61^. We found MBP in both soluble and histone-bound fractions of the MG nuclei, raising the possibility that it could directly or indirectly regulate MG gene expression. Analysis of MBP binding motifs (e.g., cut and run analysis) might provide insight into the immunopathogenesis of MS. Further work is also needed to determine whether MBP-delivered RNAs are translated or whether they modulate intrinsic MG inflammatory response programs. MG are relatively refractory to conventional gene delivery methods such as lipofection and viral transduction, representing barriers for experimental and therapeutic manipulation. We used MBP to deliver small interfering RNAs (siRNAs) to facilitate knockdown of the *Tlr4* and mitigate LPS-induced inflammation, as reflected by reduced CXCL10, IL6 and TNFA expression at the mRNA and cytokine levels. These findings have relevance to a recent study showing that maladaptive MG activation promotes CXCL10-mediated recruitment of CD8+ T cells in ageing-related white matter degeneration ^111^. The demonstration that MBP can deliver mRNA into MG nuclei suggests a potential platform for targeted transcriptome modulation and genome editing in MG.

## Author Contributions

G.L. and D.H.R conceptualized the project and designed the experiments.

G.L, A.M.V, S.V., O.D.F. Jr, M.P., J.H.S, D.Y., Q.Q.Z, G.J., J.A.S performed the experiments and analysed the data. J.Z.Y.X, S.V. conducted the data analysis.

R.T.K, D.S., A. B., M.S, H.W., A.L, H.B, J.J, L.R.M, J.S., H.M helped interpreted data and contributed to the discussion.

G.L., S.V., A.M.V and G.C.G. arranged the figures.

G.L., A.M.V, S.V. G.C.G, DHR wrote the paper.

All authors helped write and/or edit the manuscript.

## Funding

This work was supported by funding from the European Research Council Advanced Grant (789054 to D.H.R), Dr. Miriam and Sheldon G. Adelson Medical Research Foundation (to D.H.R. and D.P.S), the Wellcome Trust (to D.H.R), MS Society Centre of Excellence award (132 to R.T.K.), MRC grant MC-A025-5PK71 (M.S.), Cure Alzheimer’s Fund (D.P.S.) and the NIH [NINDS (F31 NS139677 J.M.S.), NINDS (R01 NS117533 D.P.S); NIMH-R01MH113743 (D.P.S.), NIA-RF1AG068281 (D.P.S.), NINDS PO1 NS083513 (D.H.R.)]. We further acknowledge an Early Career Fellowship (320224/Z/24/Z; S.V.) and centre award (203151/Z/16/Z, 203151/A/16/Z); the UKRI (programme grant MR/Y014537/1; R.T.K., O.d.F.) and a centre award (MC_PC_17230); MS Society Centre of Excellence award (132; R.T.K., O.d.F., S.V., M.P., B.V.V.); MS Society Early Career Fellowship (207; O.d.F.). This work was supported by the NIHR Cambridge Biomedical Research Centre (NIHR203312); views expressed are those of the authors and not necessarily those of the NIHR or the Department of Health and Social Care.

## Competing interests

The authors declare no competing interests.

## Data Availability Statement

The datasets generated during this study are available from the corresponding author upon reasonable request. Source data are provided with this paper. All sequencing data generated in this study have been deposited in the Gene Expression Omnibus (GEO) under accession number GSE330818. Additional data supporting the findings of this study are available from the corresponding author upon reasonable request.

## AI Use Declaration

The authors declare that they have not use generative AI in this manuscript.

## Acknowledgements

The authors thank members of the Rowitch, Káradóttir and Jones group laboratories for many useful discussions and helpful comments. We gratefully acknowledge Dr. Darran Clements and the Advanced Imaging Facility, University of Cambridge for their support & assistance in this work. The authors thank the Mass Spectrometry Facility from the Department of Pharmaceutical Chemistry, UCSF for their technical assistance, and Dr. Antonio García for his help on optimizations of the figure schematics.

## Methods

### Animals

Experiments were performed in accordance with the EU guidelines for the care and use of laboratory animals, and with the guidelines of the UK Animals (Scientific Procedures) Act 1986 and subsequent amendments. Use of animals in this project was approved by the Animal Welfare and Ethical Review Body for the University of Cambridge and carried out under the terms of UK Home Office Licenses (license number PP4690795). All animals were maintained under a 12 h–12 h light–dark cycle with food and water supplied ad libitum, in individually ventilated cages at 20–24 °C and 45–65% relative humidity. All surgeries were performed aseptically under isoflurane anaesthesia, with standard pre- and post-operative analgesia and care.

### Human Tissue License

Patient neurosurgical surplus tissue was obtained with informed consent under UK Health Research Authority permissions REC 18/WM/0094 (Principal Investigator Harry Bulstrode) and IRAS 238683, following all regulations and processed immediately following resection. Human tissue underwent primary CD11B+ isolation in CL2 primary human tissue culture conditions for dissociated cell adherent culture.

### Myelin purification and fluorescent labelling

Human and mouse Myelin was purified based on a modified protocol from Jahn et al. (2012), with reference to Thakurela et al. (2016). All steps were performed at 4 °C where possible to minimize RNA and protein degradation. Brain tissue was Dounce-homogenized in DEPC-treated 0.32 M sucrose (Sigma-Aldrich, S0389) supplemented with HALT protease inhibitor (Thermo Fisher Scientific, 10516495). A total of 6 ml of homogenate was carefully layered onto 6 ml of 0.85 M sucrose in 14 ml open-top polypropylene tubes (Beckman Coulter, C14303) pre-treated with RNaseZap (Thermo Fisher Scientific, R2020). Samples were subjected to ultracentrifugation at 75,000g for 35 min at 4 °C using an Optima XPN-80 ultracentrifuge (Beckman Coulter) equipped with an SW40Ti rotor. Following centrifugation, three distinct layers were observed and the interphase was collected. This sucrose gradient separation step was repeated once to further reduce cytosolic contamination. The interphase fraction was then washed in 1× PBS (Thermo Fisher Scientific, 10010023) and centrifuged at 12,500g for 15 min; this wash step was repeated three times. The final pellet, corresponding to a crude myelin debris fraction, was collected and used for subsequent myelin-feeding and nuclear uptake assays. As an alternative purification step, the interphase fraction was subjected to hypotonic treatment to reduce contaminating proteins. Briefly, the interphase was resuspended in ice-cold DEPC-treated water (Sigma-Aldrich, D5758) and incubated for 1 min at 4 °C to induce osmotic shock. Samples were then brought to 10 ml with water and centrifuged at 12,500g for 15 min. The supernatant was removed and the osmotic shock step was repeated once. The resulting pellet was resuspended in 0.35 M sucrose (1 ml) and adjusted to a final volume of 6 ml, then layered onto 0.85 M sucrose and subjected to ultracentrifugation at 75,000g for 35 min at 4 °C to re-establish density separation. The interphase fraction was collected, washed three times in 1× PBS and centrifuged at 12,500g for 15 min. The final pellet was collected as a purified myelin debris fraction. This purification step reduces cytoplasmic and neuronal contaminants; however, it may also alter the native composition of myelin. Hypotonic exposure can promote dissociation of membrane-associated proteins and loss of soluble components, particularly from non-compact myelin regions. Therefore, although certain proteins may be retained in the pellet following high-speed centrifugation, their structural association with myelin may be disrupted relative to in vivo conditions.

Total protein concentration of myelin preparations was determined using the Qubit protein assay (Thermo Fisher Scientific, Q33212). For fluorescent labelling, myelin was diluted to 1 mg ml⁻¹ in 0.1 M sodium bicarbonate buffer (pH 8.3; Sigma-Aldrich, S5761) and incubated with pHrodo Red succinimidyl ester (Thermo Fisher Scientific, P36600) at the manufacturer’s recommended working concentration for 1 h at room temperature. Labelled samples were centrifuged at 4 °C to remove excess dye, washed twice and resuspended in sterile 1× PBS prior to use. In parallel, native or recombinant myelin proteins (commercially sourced: Abcam, ab64311, ab43614, ab290085) were used in the study to investigate nuclear localization. Protein solutions first undergone buffer-exchange and were concentrated to 1 mg ml⁻¹ in 0.1 M sodium bicarbonate buffer (∼pH 8.3; Sigma-Aldrich, S5761), using 3 kDa molecular weight cut-off centrifugal filters (Pierce Protein Concentrators PES; Thermo Fisher Scientific, 88512). The setup was then incubated with pHrodo Red succinimidyl ester (Thermo Fisher Scientific, P36600) for 1 h at room temperature.

### Primary microglia isolation and culture

Primary microglia were isolated from postnatal day 8 (P8) rat pups (Charles River Laboratories; typically 6–24 heads per preparation; protocol described for 6 heads). Human microglia and myelin samples were obtained from neurosurgical surplus tissue with informed consent under UK Health Research Authority approval (REC 18/WM/0094, IRAS 238683; Principal Investigator H. Bulstrode), in accordance with the Human Tissue Act 2004 and relevant guidelines. Tissue was processed immediately following resection. Brains were dissected under sterile conditions and meninges removed. Tissue was mechanically dissociated using a scalpel and homogenized in cold HBSS (without Ca²⁺/Mg²⁺), followed by centrifugation at 100g for 3 min at room temperature. The pellet was enzymatically digested with papain (10–20 U ml⁻¹; Worthington/Lorne LS003126) for 20 min at 37 °C with gentle agitation. Samples were centrifuged at 300g for 5 min and resuspended in cold neutralization buffer (Hibernate A supplemented with B27, 1 mM sodium pyruvate and 1 µg ml⁻¹ DNase I). Cells were mechanically dissociated by serial trituration (10× with 10 ml, 10× with 5 ml and 10× using a fire-polished pipette), filtered through a 70 µm cell strainer and layered onto Percoll (final concentration 22.5%). Following centrifugation at 800g for 20 min at room temperature (no brake), the cell-containing fraction was collected, diluted in HBSS and centrifuged at 300g for 5 min. Red blood cells were lysed using RBC lysis buffer (Thermo Fisher Scientific, J62990) for 1.5 min at room temperature, diluted with HBSS and centrifuged again at 300g for 5 min. Cells were resuspended in MACS wash buffer (PBS containing 0.5% BSA, 2 mM EDTA, 1 mM sodium pyruvate and 50 µg ml⁻¹ insulin) and incubated with CD11b/c MicroBeads (Miltenyi Biotec, 130-105-643, 130-110-614; 20 µl per 10⁷ cells) for 15 min at 4 °C. After washing, cells were applied to LS columns (Miltenyi Biotec, 130-042-401) mounted on a MACS separator and washed three times with buffer. CD11b⁺ cells were eluted following column removal and gentle flushing. Isolated microglia were counted and plated at a density of 3 × 10⁴ to 1 × 10⁵ cells cm⁻². For human microglia isolation, the same protocol was used with anti-human CD11b antibody (Miltenyi Biotec, APC-Vio770 REAfinity) at 1:50 dilution. Human tissue processing and culture were conducted under CL2 conditions. Microglia were cultured in defined microglia culture medium consisting of DMEM/F12 supplemented with N2, 1 mM sodium pyruvate, 50 µg ml⁻¹ insulin, 3 mg ml⁻¹ N-acetylcysteine, 1.5 µg ml⁻¹ cholesterol, macrophage colony-stimulating factor (mCSF; 20 ng ml⁻¹ for rat, 100 ng ml⁻¹ for human) and transforming growth factor-β (TGFβ; 2 ng ml⁻¹ for rat, 5 ng ml⁻¹ for human), with optional penicillin–streptomycin. Feeding experiments were performed the day after plating to allow cell adherence and recovery.

### RNA Synthesis and Purification

Mouse Mobp (NM_001039365) in pcDNA3.1-3xFLAG (NovoPro, 709264-1), pGEMHE-NLS-mEGFP (Addgene, Plasmid #105527-sequence-201844), pmRVac-rSt18 HA (VectorBuilder, Custom) were used as vector templates for RNA synthesis. PCR Amplification of DNA Template was conducted with Kappa Roche Diagnostics KAPA HiFi HotStart ReadyMix PCR Kit (Roche, 50-196-5217), with the following primers: Mobp PCR Primers: Forward Primer (CMV-F): 5’-CGCAAATGGGCGGTAGGCGTG-3’, Reverse Primer (F1Ori-R): 5’-AGGGAAGAAAGCGAAAGGAG-3’. EGFP PCR Primers: Forward Primer (CMV-F): 5’-CGCAAATGGGCGGTAGGCGTG-3’, Reverse Primer (M13 Reverse): 5’-CAGGAAACAGCTATGAC-3’. ST18 PCR Primers: Forward Primer (CMV-F): 5’-AGCGAGTCAGTGAGCGAG-3’, Reverse Primer (M13 Reverse): 5’-GGGTTATTGTCTCATGAGCG-3’

DNA templates were purified using spin columns (Qiagen, 27104), and size and yield were assessed by agarose gel electrophoresis (Thermo Fisher Scientific, G401002) and NanoDrop spectrophotometry. In vitro transcription was performed using the HiScribe T7 ARCA mRNA Kit (New England Biolabs, E2065S) according to the manufacturer’s instructions. Following transcription, RNA was purified by phenol–chloroform extraction (TRIzol reagent, Invitrogen, 15596026; chloroform, Sigma-Aldrich, 372978) and resuspended in RNase-free water. For biotin labelling, RNA was modified using the Pierce RNA 3′ End Desthiobiotinylation Kit (Thermo Fisher Scientific, 20163). Alternatively, DIG-labelled RNA probes were generated using the DIG RNA Labelling Kit (T7) (Sigma-Aldrich, 11175025910). For DIG-labelled RNA, phenol–chloroform extraction was avoided to prevent loss of DIG-labelled transcripts to the organic phase; instead, purification was performed using chloroform–isoamyl alcohol extraction.

### Preparation of myelin/ protein–RNA complexes and microglial feeding assays

Myelin was applied to cultured microglia at 1 or 10 μg ml⁻¹ unless otherwise specified. For pHrodo Red–labelled protein assays, MBP (Abcam, ab64311, ab43614) and related myelin proteins were typically applied at 1 μg ml⁻¹ to enable sensitive detection. For immunocytochemistry (ICC)-based detection, MBP was applied at 10–200 μg ml⁻¹. For *in vitro* RNA trafficking experiments, MBP was applied at 50–200 μg ml⁻¹ in combination with DIG-labelled RNA (10 μg ml⁻¹), for robust nuclear detection. Protein–RNA complexes were prepared at defined mass ratios (10:1–50:1 for mRNA delivery, 10:1–100:1 for siRNA, and 10:1–25:1 for TLR stimulation assays). MBP was added to RNA and incubated for 10 min at room temperature to allow complex formation prior to addition to culture medium. For delivery of unlabelled RNA, mRNA was applied at 500 ng ml⁻¹ and detected by single-molecule FISH (smFISH). For myelin–RNA feeding experiments, myelin was applied at 10–100 μg ml⁻¹ with corresponding RNA concentrations as indicated above. For siRNA delivery, flocculation was minimized by gradual complex formation. siRNA (Thermo Fisher Scientific, 4390771, 1320001, s14194) was used either undiluted or diluted in 30 μl microglia culture medium (∼150 mM ionic strength). Protein was added incrementally in small volumes to the RNA solution with gentle mixing to prevent local charge neutralization and promote stable complex formation.

### Brain injection of MBP–Mobp protein–RNA complexes and tissue processing

MBP–Mobp complexes were prepared and incubated for 10 mins at room temperature to allow complex formation, immediately prior to injection. DIG-labelled Mobp RNA (500 ng) was incubated with native or recombinant MBP (Abcam; ab64311 and ab43614), pHrodo Red-labelled MBP alternatives, or MAG (Abcam, ab290085), to yield final doses of 0.2, 1, or 3 μg protein per 500 ng RNA. Male C57BL/6 mice (P8, 3–4 months old respectively) were anaesthetized and subjected to bilateral stereotaxic injections into the corpus callosum using a 10 μl Hamilton syringe. MBP–Mobp complexes (2 μl per hemisphere) were delivered at a rate of 0.25 μl min⁻¹ at the following coordinates relative to bregma: −2.5 mm anterior–posterior (AP), ±0.8 mm mediolateral (ML), and −1.2 mm dorsal–ventral (DV). For paired hemispheric injections, experimental and control conditions were administered into opposite hemispheres of the same animal. For gradient-dependent trafficking analyses, the following conditions were used:

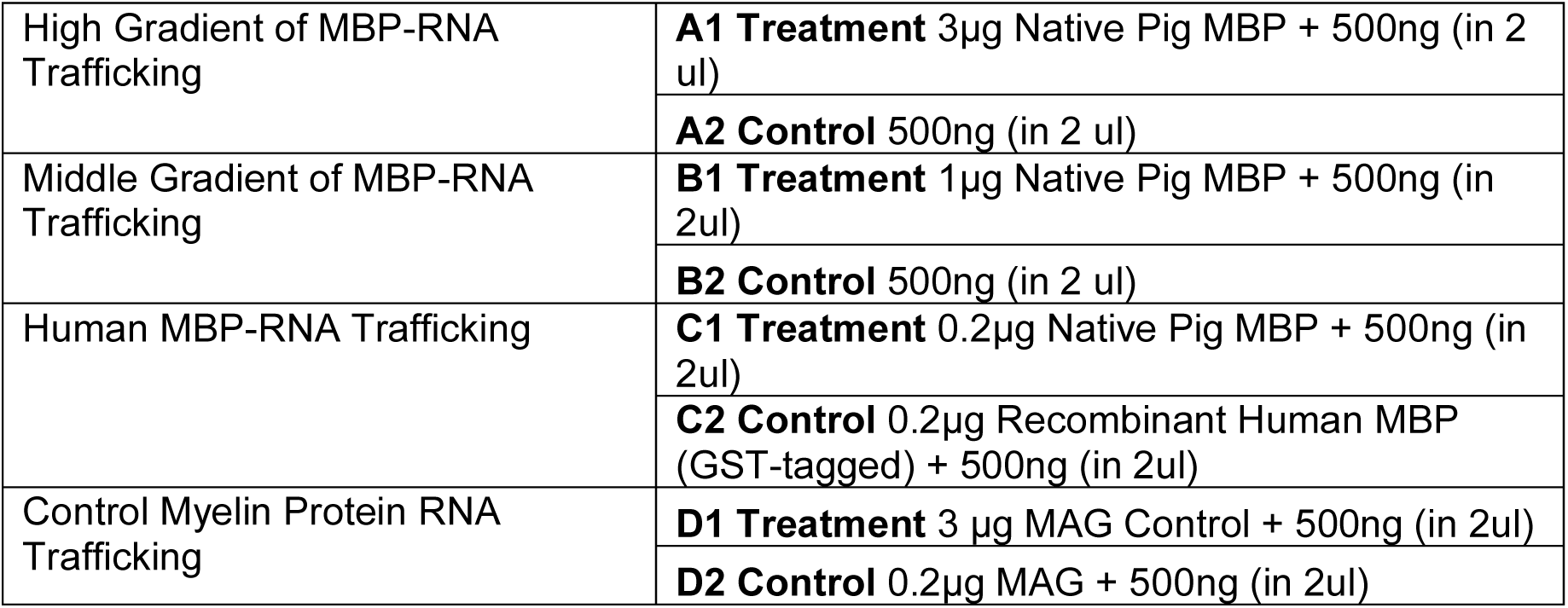

|  |  |
| --- | --- |
| High Gradient of MBP-RNA Trafficking | **A1 Treatment** 3μg Native Pig MBP + 500ng (in 2 μl) |
| | **A2 Control** 500ng (in 2 μl) |
| Middle Gradient of MBP-RNA Trafficking | **B1 Treatment** 1μg Native Pig MBP + 500ng (in 2μl) |
| | **B2 Control** 500ng (in 2 μl) |
| Human MBP-RNA Trafficking | **C1 Treatment** 0.2μg Native Pig MBP + 500ng (in 2μl) |
| | **C2 Control** 0.2μg Recombinant Human MBP (GST-tagged) + 500ng (in 2μl) |
| Control Myelin Protein RNA Trafficking | **D1 Treatment** 3 μg MAG Control + 500ng (in 2μl) |
| | **D2 Control** 0.2μg MAG + 500ng (in 2μl) |

At 3 days post-injection, animals were transcardially perfused with heparinized PBS (pH 7.4), followed by 4% paraformaldehyde (PFA) in PBS. Brains were dissected and post-fixed in 4% PFA for 1 h at 4 °C, then cryoprotected overnight in 30% sucrose (Sigma-Aldrich, S0389) in PBS at 4 °C. Tissues were subsequently equilibrated in 30% sucrose/OCT for 30 min at room temperature, followed by OCT alone (Thermo Fisher Scientific, 15212776) for 30 min. Brains were embedded on dry ice and stored at −80 °C or sectioned immediately. Coronal cryosections (30–50 μm) were cut using a CM3050 S cryostat (Leica Microsystems) and mounted onto slides or stored at −80 °C until further immunofluorescence analysis.

### Immunocytochemistry and immunohistochemistry

Cultured cells were washed twice with 1× PBS and fixed in 4% paraformaldehyde (PFA; pH 7.4, supplemented with Ca²⁺ and Mg²⁺) for 15 min at room temperature. Cells were then washed three times with PBS (10 min each) with gentle agitation. For immunocytochemistry (ICC), cells were blocked in 5% normal serum (matched to the secondary antibody host species; normal donkey serum (Abcam, ab7475) or goat serum (Sigma-Aldrich, G6767)) supplemented with 0.05% Triton X-100 (Sigma-Aldrich, T8787) and 0.05% digitonin (Abcam, ab300022). Primary antibodies were diluted in blocking buffer (1:250; ∼2.5 μg ml⁻¹) and incubated overnight at 4 °C. Following incubation, cells were washed three times with PBS (10 min each) and incubated with secondary antibodies (1:500; ∼5 μg ml⁻¹) in blocking buffer for 1 h at room temperature. Nuclear counterstaining was performed using Hoechst (10 μg ml⁻¹; 1:5000) during the final 10 min of secondary antibody incubation. Cells were washed three times with PBS prior to imaging. For immunohistochemistry (IHC), brain sections were subjected to antigen retrieval in citrate buffer (10 min at 80 °C) prior to blocking. Sections were blocked in 5% normal donkey serum and 3% BSA containing 0.5% Triton X-100 for 1 h at room temperature. Primary antibodies were applied at 1:300 for 1 h at room temperature. Sections were washed three times in PBST (0.5% Triton X-100 in PBS), followed by incubation with secondary antibodies (1:1000) for 1 h at room temperature. After three washes in PBST, sections were rinsed in PBS and counterstained with Hoechst, followed by additional washes in PBST. Sections were mounted using Fluoromount-G (Invitrogen, 15586276). Reagents, antibodies and plasmids are listed in Supplementary Table 3.

### Multiplex in situ RNA hybridization (smFISH)

Microglia cultured on slides were fixed in 4% paraformaldehyde (PFA) and incubated with hydrogen peroxide (Sigma-Aldrich, H1009) for 10 min at room temperature. Slides were washed three times in water (5 min each) and subjected to target retrieval in ER2 buffer (Leica, AR9640) for 3 min at 95 °C. Following retrieval, slides were washed in water (three times, 5 min each) and dehydrated in 100% ethanol (Sigma-Aldrich, 51976). After air drying, a hydrophobic barrier was drawn around samples using a PAP pen. Sections were then treated with Protease IV for 30 min at 40 °C, followed by hybridization with target probes for 2 h at room temperature. Signal amplification was performed sequentially using AMP1 (30 min), AMP2 (30 min) and AMP3 (15 min) (Leica RNAscope Multiplex Fluorescent Reagent Kit; DS9790). Slides were then incubated with HRP-conjugated probes (HRP-C1, HRP-C2 and HRP-C3; 15 min each), with signal development using Opal fluorophores (Akoya Biosciences) in TSA buffer (30 min each). HRP blocking steps (15 min) were performed between each channel to prevent cross-reactivity. Following RNA detection, slides were counterstained with DAPI and processed for immunofluorescence using primary and secondary antibodies as described above.

### Microscopy and image acquisition

Fluorescence imaging and live-cell imaging were performed using a confocal laser scanning microscope (LSM 980; ZEISS) equipped with Airyscan. Representative images were acquired using a 63× oil immersion objective (NA 1.4), with a pixel size of 2924 × 2924, 16 -bit depth and zoom factor of 1.3. High-content tile scans for quantitative analysis were acquired using an Operetta CLS system (PerkinElmer). Perkin Elmer scanning confocal microscope and Harmony automated quantification were conducted to determine the nuclear pHrodo red^+^ area/ object ratio (pHrodo red^+^ Iba1^+^ DAPI^+^ area in μm^2^/ pHrodo red^+^ MG objects) normalized to positive myelin-fed control. For imaging of IHC-stained brain sections, a Leica Stellaris 8 confocal microscope was used with a 63× oil immersion objective (NA 1.4). Imaging parameters were as follows: scanning speed 600, pixel size 45.07 × 45.07 nm, line averaging 1, z-step size 0.15 μm, bit depth 12 and pinhole size 1 Airy unit. All images were acquired using identical laser power settings and hybrid (HyD) detectors in photon-counting mode. Six images were acquired per animal (three per injection site). Back-projected pinhole was calculated per fluorophore used (DAPI, Alexa488, Alexa568, Alexa647) using the following formula:

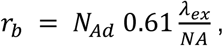

Where r_b_ is the back-projected pinhole radius, N_Ad_ is the number of Airy disks (provided by the SVI website). The values for the back-projected pinhole were 176.5, 212.6, 240.5, 281.9 respectively. Huygens Professional (SVI) was used to deconvolve the high-resolution images. All channels were deconvolved with the Classic Maximum Likelihood Estimation (CMLE) algorithm using default settings apart from the following:

| Channel | DAPI | Alexa488 | Alexa568 | Alexa647 |
| --- | --- | --- | --- | --- |
| Nr of iterations | 50 | 100 | 100 | 100 |
| Acuity | 0 | 0 | 10 | 10 |

### Image analysis and quantification: nuclear segmentation and DIG particle analysis

Deconvolved high-resolution micrographs were imported into Imaris 10.2 (Bitplane). The Surfaces tool was used to generate a surface rendering of DAPI positive nuclei using automated thresholding. Then, the Surface was used to mask the DIG channel. The DAPI channel and the masked DIG channel were used in the Cells tool (as cell+vesicles) to segment individual nuclei. Since each brain had a control and a treatment side, the control side was used to threshold the DIG signal based on the maximum intensity of the masked DIG channel, and the same threshold was applied to the treated side. Object-object statistics were exported, and numbers of DIG particles were analysed.

### Subcellular Fractionation: Nuclear Isolation for Transcriptomics vs Proteomic Analysis

Nuclear Isolation for transcriptomic analysis was done with 10x protocol ‘Nuclei Isolation from Cell Suspensions & Tissues for Single Cell RNA Sequencing’ CG000124. Subcellular Fraction for proteomics analysis was done with either NE-PER™ Nuclear and Cytoplasmic Extraction Reagents (Thermo Fisher, 78833), or Subcellular Protein Fractionation Kit for Cultured Cells (Thermo Fisher, 78840).

### RNA extraction, qPCR and RNA-seq library preparation

Cells or subcellular fractions were washed with 1× ice-cold PBS and lysed in TRIzol reagent (Invitrogen, 15596026) for >5 min at room temperature to ensure complete dissociation of nucleoprotein complexes. Chloroform (0.2 ml per 1 ml TRIzol; Sigma-Aldrich, 372978) was added. mRNA was subsequently purified using the PureLink RNA Mini Kit with on-column DNase treatment (Thermo Fisher Scientific, 12183018), following manufacturer’s instructions. RNA integrity and concentration were assessed using the RNA ScreenTape system (reagent: Agilent, #5067-5577, RNA SceenTape: Agilent, 5067-5576, TapeStation: 4200 TapeStation System, Agilent) and NanoDrop spectrophotometry. Only high-quality RNA (RNA integrity number (RIN) > 7) was used for downstream applications. For reverse transcription, RNA was processed using the ZymoScript RT PreMix Kit (Zymo Research, R3012). For RNA sequencing, libraries were prepared using the QIAseq UPXome RNA Lib Kit HMR (poly(A) selection; Qiagen, HB3095) and Qiagen Upxome Low Input Kit (334705), according to the manufacturer’s protocol. Prepared RNA-seq libraries were subjected to bulk sequencing on an Illumina NovaSeq 6000 platform using the NovaSeq SP paired-end 150 bp configuration (PE150) (Illumina, San Diego, CA, USA).

Raw paired-end FASTQ files underwent initial quality assessment with fastqc. Illumina adapter sequences and low-quality bases were then trimmed using trimmomatic (v0.39) (RRID:SCR_011848), ensuring that only high-quality reads were retained for downstream analysis.

### RNA–protein pulldown

Subcellular fractions from DIG–RNA-fed microglia were prepared in lysis buffer and incubated with anti-digoxigenin magnetic beads (25–50 μl per sample; Thermo Fisher Scientific) in 1.5 ml microcentrifuge tubes, following a protocol adapted from the Pierce Magnetic RNA–Protein Pull-Down Kit (Thermo Fisher Scientific, 20164). Bead volumes were adjusted according to input material (for example, 50 μl beads bind ∼3 pmol RNA (∼1.23 μg)). Samples were incubated overnight at 4 °C with rotation to allow DIG-specific binding. Tubes were then placed on a magnetic rack (DynaMag-2; Thermo Fisher Scientific, 12321D) to separate bead-bound complexes from the supernatant. The supernatant fraction was retained as input (total nuclear proteome). Beads were washed three times with wash buffer (100 μl per wash) to remove non-specifically bound proteins, followed by two additional washes in 20 mM Tris (pH 7.5; Sigma-Aldrich, T5912) to reduce background for downstream mass spectrometry analysis. Beads were resuspended by gentle pipetting or vortexing between washes. After the final wash, beads were collected using the magnetic rack and stored at −80 °C until further analysis. For elution of RNA–protein complexes, beads were incubated with elution buffer (Thermo Fisher Scientific, 20164) for 15–30 min at 37 °C with agitation. Eluted samples were mixed with tricine SDS sample buffer (Thermo Fisher Scientific, LC1676), heated at 75 °C for 5 min and resolved by SDS–PAGE for downstream analysis (for example, silver staining).

### RNA–protein binding assay and EMSA

Biodyne B nylon membranes (Thermo Fisher Scientific, 77016) were pre-wetted in 0.5× TBE buffer (Alfa Aesar, J62788) to ensure uniform hydration. Gradients of MBP or control proteins were incubated with RNA probes in 1× REMSA binding buffer (LightShift Chemiluminescent RNA EMSA Kit; Thermo Fisher Scientific, 20158) for 5 min at room temperature. Samples were applied to the membrane using a Bio-Dot microfiltration system (Bio-Rad, 1703938) under gentle vacuum. The membrane was maintained under suction for ∼15 min to allow binding prior to complete drying. Following application, membranes were UV crosslinked for 50 s. Biotin-labelled RNA was detected using the LightShift Chemiluminescent Nucleic Acid Detection Module (Thermo Fisher Scientific, 89880) according to the manufacturer’s instructions. To assess RNA–protein complex stability (“melt-off”), membranes were sealed and incubated sequentially in water baths at increasing temperatures for 1 min per condition, followed by imaging after each step. Electrophoretic mobility shift assays (EMSA) were performed using the LightShift Chemiluminescent RNA EMSA Kit and associated optimization and control reagents (Thermo Fisher Scientific, 20158X, 89880) according to the manufacturer’s protocol.

### Western blot analysis

Protein samples were quantified and prepared for western blotting in tricine SDS sample buffer (Thermo Fisher Scientific, LC1676) supplemented with reducing agent (NuPAGE, Thermo Fisher Scientific). Samples were adjusted to a final volume of 100 μl with water, and 15–30 μg total protein was loaded per lane. Samples were denatured at 75 °C for 2 min and resolved on 10–20% Tricine SDS–PAGE gels (Thermo Fisher Scientific, EC6625BOX) at 90 V for 20 min followed by 120 V for 100 min. Proteins were transferred onto PVDF membranes (Thermo Fisher Scientific, 88518) at 15 V for 90 min. Membranes were blocked in 3% BSA (Sigma-Aldrich, A8327) for 1 h at room temperature and incubated with primary antibodies (1:1000) overnight at 4 °C. After washing in TBST (Promega, H5151), membranes were incubated with secondary antibodies and washed as above. Signals were detected using an Odyssey LI-COR imaging system.

### TMT-based mass spectrometry analysis

Proteins were recovered from microglial subcellular fractions (obtained from 2 million cells as described above), by aggregation on carboxylate-coated magnetic beads in organic solvent, essentially as described (Hughes CS, Moggridge S, Müller T, Sorensen PH, Morin GB, Krijgsveld J. Single-pot, solid-phase-enhanced sample preparation for proteomics experiments. Nat Protoc. 2019 Jan;14(1):68-85. doi: 10.1038/s41596-018-0082-x. PMID: 30464214.). Samples in the different fractionation buffers (50 to 100 ul) were added 530 μg of washed Sera-Mag Speed Beads (Thermo Fisher CAT# 09-981-121) and then added an equal volume of ethanol, and incubated for 8 minutes at RT. After that, samples were placed in a magnetic rack and the supernatant removed. Beads were washed twice with 200 μl 80% Ethanol, then supernatant removed and beads resuspended in 20 μl of 10 mM Tris(2-carboxyethyl)phosphine (TCEP) plus 20 mM triethylammonium bicarbonate (TEAB) pH 8.5, and incubated for 30 minutes at 56°C. After this, iodoacetamide was added to a final concentration of 20 mM and samples incubated at room temperature for 60 additional minutes. 0.5 μg of sequencing grade trypsin (Promega) was added to each sample and incubated at 37°C overnight. Supernatants were recovered, and beads were digested again using 0.5 μg trypsin in 9 μl 20 mM TEAB for 4 hours at 37°C. Peptides from both consecutive digestions were combined and subjected to TMT labelling. For TMT labelling, samples were labelled according to the TMT 16plex kit instructions (ThermoFisher Scientific), with minor modifications. Briefly, TMT reagents were resuspended in acetonitrile at 25 μg/μl, and 4 μl of this solution were added to the individual samples to be labelled. After incubating for 1 hour at 22°C, reactions were quenched by adding 4 μl 5% hydroxylamine and incubated for an additional 15 minutes. After that, the labelling reactions were combined and diluted then all samples were combined adding them over 20 ml 0.1% formic acid, and desalted using a C18 SepPak cartridge (Waters). The Sep Pak eluate was dried in preparation for chromatographic fractionation on an AKTA purifier system utilizing a Phenomenex Gemini 5u C18 110A 100mm x 1 mm column, operating at a flow rate of 0.080 ml/min. Buffer A consisted of 20 mM ammonium formate (pH 10) , and buffer B consisted of 20 mM ammonium formate in 90% acetonitrile (pH 10). Gradient details were as follows: 1 % to 30% B in 120 min, 30% B to 70% B in 20 min, 70% B down to 1% B in 20 min. Peptide-containing fractions were collected, evaporated and resuspended in 0.1% formic for LCMSMS analysis. Fractions (9 to 13 depending on the experiment) of labelled peptide digests were subjected to chromatographic separation using a 2 μm, 75μm ID x 50 cm PepMap RSLC C18 EasySpray column (Thermo Scientific). 3-hour MeCN gradients (2–25% in 0.1% formic acid) were used to separate peptides, at a flow rate of 200 nl/min, for analysis in a Orbitrap Exploris 480 (Thermo Scientific) in positive ion mode with the following settings. MS spectra were acquired between 375 and 1500 Th with a resolution of 120000. For each MS spectrum, multiply charged ions over the selected threshold (2E4) were selected for MSMS in cycles of 3 seconds with an isolation window of 0.7 Th. Precursor ions were fragmented by HCD using stepped relative collision energies of 30, 35 and 45 in order to ensure efficient generation of sequence ions as well as TMT reporter ions. MSMS spectra were acquired in centroid mode with resolution 60000 from m/z=120. A dynamic exclusion window was applied which prevented the same m/z from being selected for 30s after its acquisition. Peak lists were generated using PAVA in-house software (Guan, S., Price, J.C., Prusiner, S.B., Ghaemmaghami, S., and Burlingame, A.L. (2011). Mol Cell Proteomics 10, M111 010728.). All generated peak lists were searched against the mouse subset of the SwissProt database (SwissProt.2019.07.31), using Protein Prospector (Clauser, K.R., Baker, P., and Burlingame, A.L. (1999). Analytical chemistry 71, 2871-2882) with the following parameters: Enzyme Specificity” was set as Trypsin, and up to two missed cleavages per peptide were allowed. Carbamidomethylation of cysteine residues, and TMT 16plex labeling of lysine residues and N-terminus of the protein were allowed as fixed modifications. N-acetylation of the N-terminus of the protein, loss of protein N-terminal methionine, pyroglutamate formation from peptide N-terminal glutamines, oxidation of methionine were allowed as variable modifications. Mass tolerance was 4 ppm in MS (after systematic error correction) and 30 ppm in MS/MS. The false positive rate was estimated by searching the data using a concatenated database which contains the original SwissProt database, as well as a version of each original entry where the sequence has been randomized. A 1% FDR was permitted at the protein and peptide level. For quantification only unique peptides were considered; peptides common to several proteins were not used. Relative quantification of peptide abundance was performed via calculation of the intensity of reporter ions corresponding to the different TMT labels, present in MS/MS spectra. Intensities were determined by Protein Prospector. For each spectrum, relative abundances in the pulldowns were calculated as ratios of each of the reporter ions intensities vs the average intensities in the appropriate reference experimental group. For peptide and then total protein relative levels, peptide spectral matches and then peptide ratios were aggregated to the peptide and protein levels using median values of the log2 ratios RNA pulldown samples were run in SDS-PAGE gels and digested in-gel with trypsin as described (J. Rosenfeld, J. Capdeveille, J.C. Guillemot and P. Ferrara. Anal. Biochem. 203 (1992), pp. 173–179) (donatello.ucsf.edu/ingel.html). The extracted digests were vacuumevaporated and resuspended in 1 μl 0.1% formic acid in water, then were subjected to chromatographic separation using a 2 μm, 75μm ID x 15 cm PepMap RSLC C18 EasySpray column (Thermo Scientific). 1 hour MeCN gradients (2–25% in 0.1% formic acid) were used to separate peptides, at a flow rate of 600 nl/min, for analysis in a Orbitrap Lumos Fusion (Thermo Scientific) in positive ion mode with the following settings. MS spectra were acquired between 375 and 1500 m/z with a resolution of 120000. For each MS spectrum, multiply charged ions over the selected threshold (2E4) were selected for MSMS in cycles of 3 seconds with an isolation window of 1.6 Th. Precursor ions were fragmented by HCD using stepped relative collision energies of 30. MSMS spectra were acquired in centroid mode with resolution 15000. Dynamic exclusion window was applied which prevented the same m/z from being selected for 30s after its acquisition.Peak lists were generated and data searched as described above excepting not allowing the TMT 16plex labelling in the searches.

### Cytokine Assay

Cytokine assays in the study employed meso scale discovery (MSD) U-Plex (custom 3-plex) mouse cytokine assay panel. Experiments were conducted by the Core Biochemical Assay Laboratory (CBAL) in Addenbrooke’s Hospital, Cambridge.

### Analysis of RNA-Seq Data

Raw paired-end FASTQ files underwent initial quality assessment with FastQC. Reads were demultiplexed using UMI-tools according to the sample barcode and UMI structure of the QIAseq UPXome library preparation workflow. Illumina adapter sequences and low-quality bases were then trimmed using Trimmomatic v0.39 (RRID:SCR_011848), ensuring that only high-quality reads were retained for downstream analysis. The cleaned reads were quantified using RSEM v1.3.3 in paired-end mode, with STAR aligner v2.7.10a used as the alignment backend within a Singularity container. For mixed-species analyses, reads were quantified against a combined mouse–rat reference generated from Rattus norvegicus Rnor_6.0 release 102 and Mus musculus GRCm39 release 104 annotations, with species-specific gene and transcript identifiers retained to distinguish mouse-derived myelin transcripts from rat microglial transcripts. RSEM generated gene- and transcript-level expected counts and normalized expression values, reported as transcripts per million (TPM). Gene-level RSEM expected counts were extracted and used as the unnormalized count-level input matrix for downstream analysis in OmicsPlayground v2.8.19. Normalization, differential expression testing, pathway enrichment analysis, unsupervised clustering and data visualization were performed within OmicsPlayground using the expected-count input matrix, with an FDR threshold of 0.05. TPM values were generated by RSEM but were not used as the primary input for OmicsPlayground downstream analyses. Quality-control reports from all steps, including FastQC, UMI-tools, Trimmomatic, STAR and RSEM, were aggregated using MultiQC to provide a comprehensive overview of data quality. Heatmaps were generated using Heatmapper. MCL clustering with an inflation parameter of 1.5 and DBSCAN clustering with epsilon = 15 were generated using STRING v12.0 (RRID:SCR_005223).

### Statistics and reproducibility

Statistical analyses were performed using GraphPad Prism (v10–11, GraphPad Software Inc, La Jolla, CA, United States) and R (RStudio). Data are presented as mean ± s.e.m. or mean ± s.d., as indicated in the figure legends. Sample sizes (n), statistical tests and *p* values are reported in the corresponding figure legends. Data distribution was assessed for normality using the Shapiro–Wilk or D’Agostino–Pearson tests, and variance homogeneity was evaluated using the Brown–Forsythe test (or Bartlett’s test, where appropriate). For comparisons between groups, one-way ANOVA, two-way ANOVA or Welch’s t-test or Welch’s ANOVA was used for normally distributed data, and the Kruskal–Wallis test was used for non-parametric data. Multiple comparisons were performed using Tukey’s, Dunnett’s or Šídák’s tests, as appropriate. Nested one-way ANOVA with Dunnett’s multiple comparisons test was used for analyses of DIG⁺ particle counts per cell and knockdown studies. Correlations were assessed using Pearson’s correlation coefficient. For nuclear localization analyses, outliers were identified using the Tukey method (1.5× interquartile range) according to a predefined criterion and excluded prior to analysis. Statistical significance is indicated in figures as follows: \**p <* 0.05, \*\**p <* 0.01, \*\*\**p <* 0.001 and \*\*\*\**p <* 0.0001.

## Data and Software availability

Figures were prepared using ImageJ 1.54f, Adobe Illustrator 28.7.4, Omero and Prism 10 and 11.

## Reporting summary

Further information on research design is available in the Nature Portfolio Reporting Summary linked to this Article.

## Reagents and resources

See supplemental table for full listing.

## Contact for reagent and resource sharing

Further information and requests for reagents may be directed to, and will be fulfilled by the corresponding author, Professor David H Rowitch.

---

**Extended Data Fig 1.**
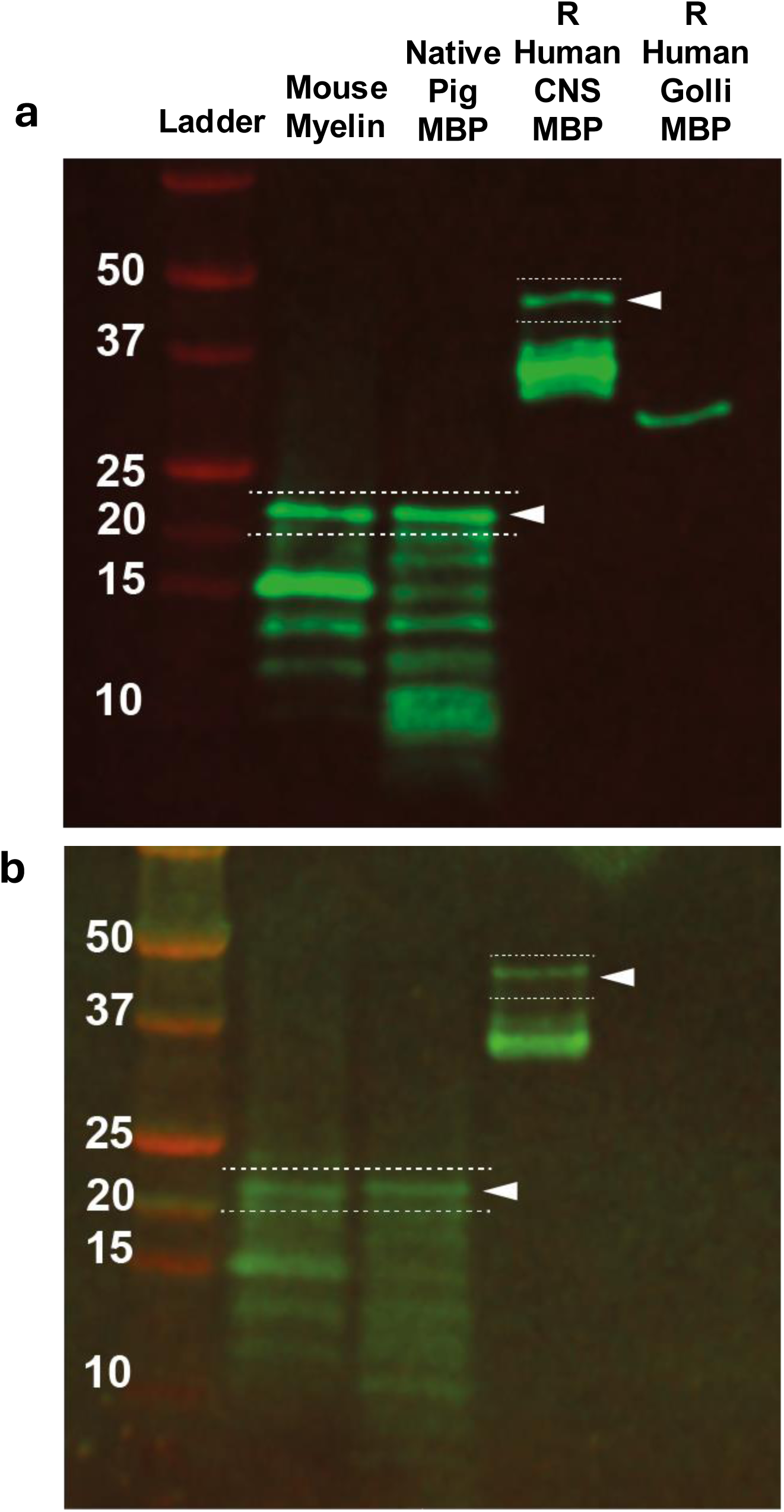
Characterization of myelin and MBP protein variants with anti-MBP antibodies. **a,** Western blot characterization of native myelin and commercially available MBP preparations. MBP isoforms were resolved and detected using an anti-MBP antibody (Cell Signaling Technology, D8X4Q, #78896). Lanes: 0, molecular weight marker; 1, mouse myelin; 2, native pig MBP (Abcam, ab64311); 3, recombinant human MBP (26kDa GST-tagged; Abcam, ab43614); 4, recombinant human Golli-MBP (5kDa His-tagged; AMSBIO, PROTP02686). The ∼21.5 kDa MBP isoform is highlighted and detected across myelin, native, and recombinant MBP samples, alongside additional MBP isoforms. **b,** Western blot characterization of native myelin and commercially available MBP preparations. MBP isoforms were resolved and detected using an anti-degraded MBP antibody (Merck/Sigma, AB5864). Similar banding patterns were observed using additional anti-MBP antibodies (Encor, MCA-7G7; Bio-Rad, MCA409S).

**Extended Data Fig 2.**
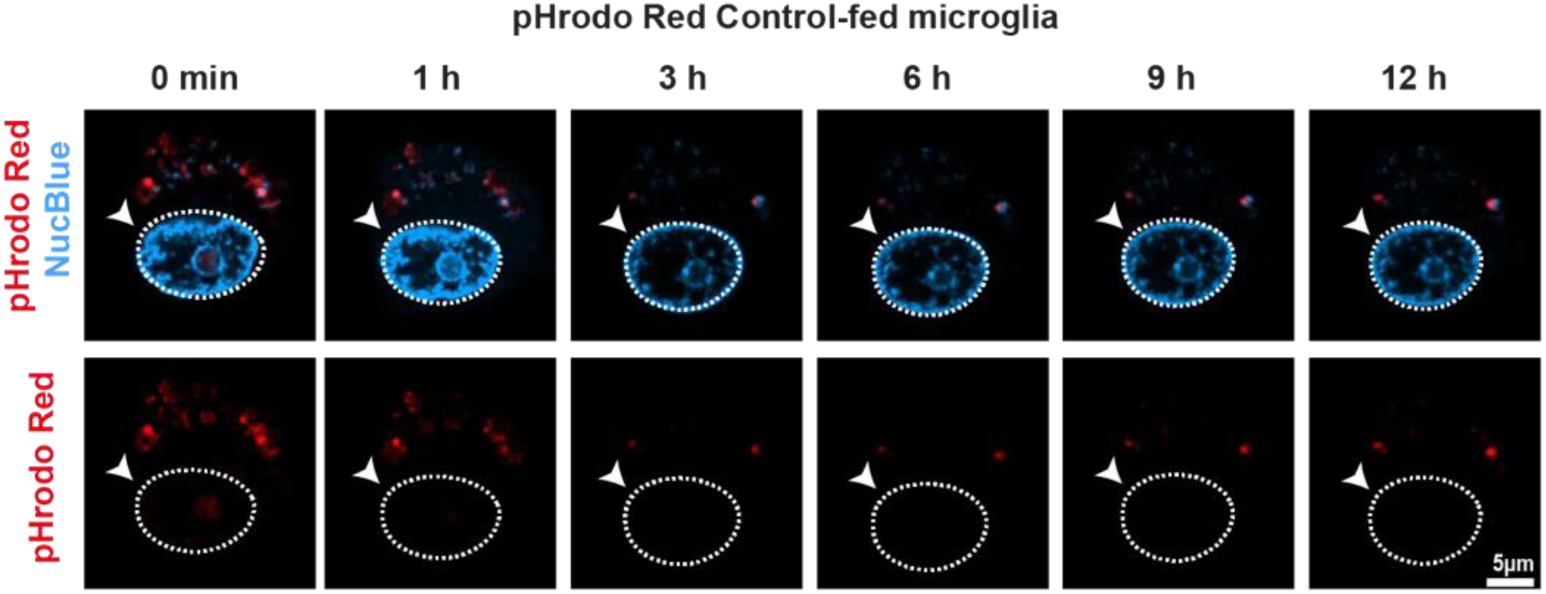
pHrodo red only does not increase in MG nucleus overtime. Representative confocal time-lapse imaging of microglia (MG) exposed to pHrodo Red–labelled control particles. pHrodo Red fluorescence (red) and nuclear staining (NucBlue^TM^, blue) are shown at the indicated time points (0–12 h). Nuclei are outlined (dashed white lines). Nuclear pHrodo Red signal remains not enriched from the nuclear compartment throughout the imaging period. Lower panels show pHrodo Red signal alone. Scale bar, 5 µm.

**Extended Data Fig 3.**
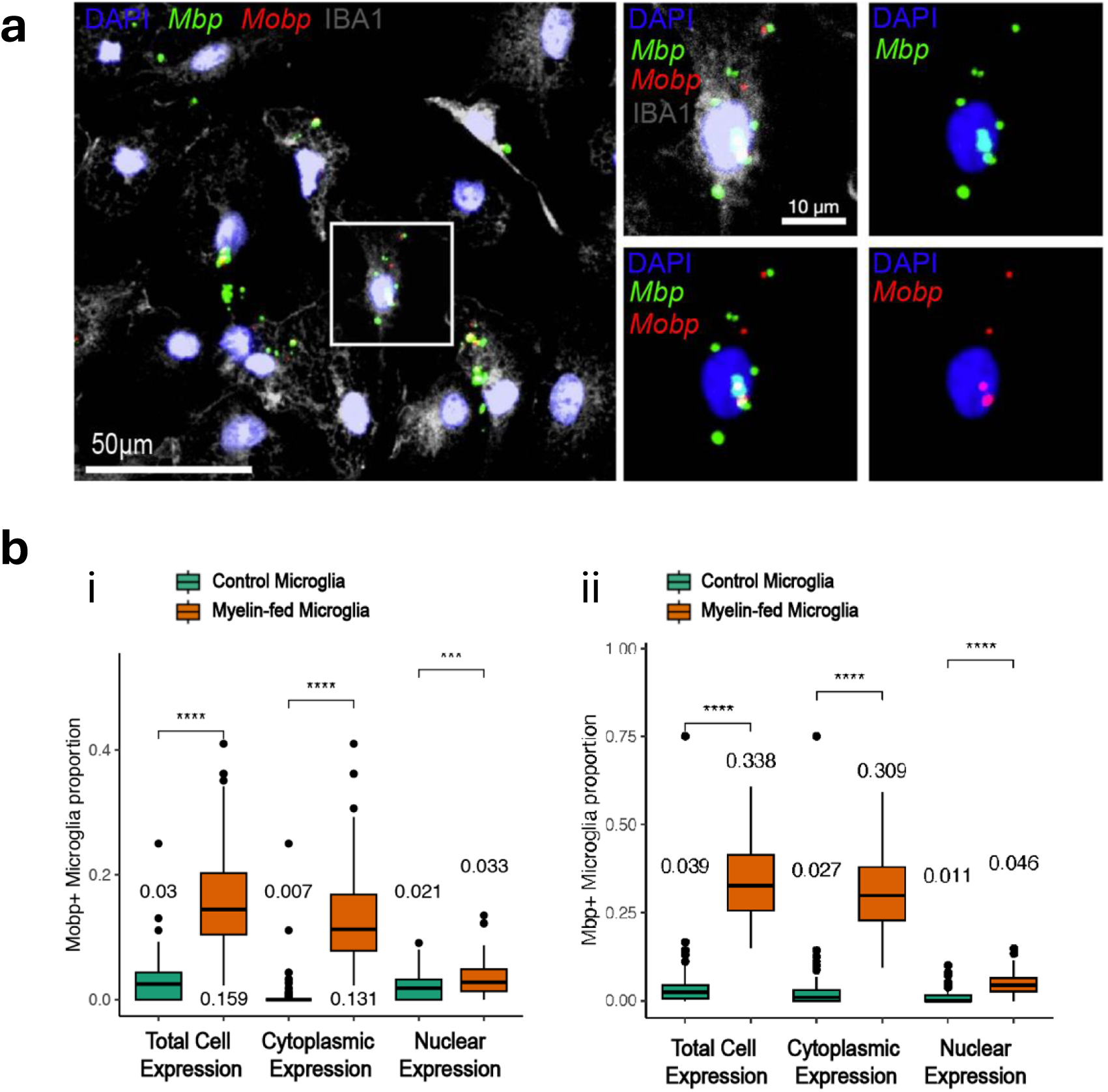
smFISH detection of major myelin transcripts in MG nucleus. **a**, Representative confocal images showing single-molecule RNA fluorescence *in situ* hybridization (smFISH) detection of *Mbp* (green) and *Mobp* (red) transcripts in IBA1⁺ microglia (grey). Nuclei are labelled with DAPI (blue). Right panels show higher-magnification views of the boxed region, with individual channels displayed separately. Scale bars, 50 µm (left) and 10 µm (right). **b**, Quantification of *Mobp* (i) and *Mbp* (ii) transcript-positive microglia in control and myelin-fed conditions. Box plots show the proportion of microglia with detectable transcripts at the whole-cell, cytoplasmic and nuclear levels. Data are presented as mean ± s.d. (*n* = 3), with individual data points representing biological replicates. Boxes indicate the median and interquartile range; whiskers denote the minimum and maximum values. Statistical analysis was performed using one-way ANOVA followed by Dunnett’s multiple comparisons test against the control condition. Adjusted *P* values are indicated (\*\*\**p* < 0.001, \*\*\*\**p* < 0.0001).

**Extended Data Fig 4.**
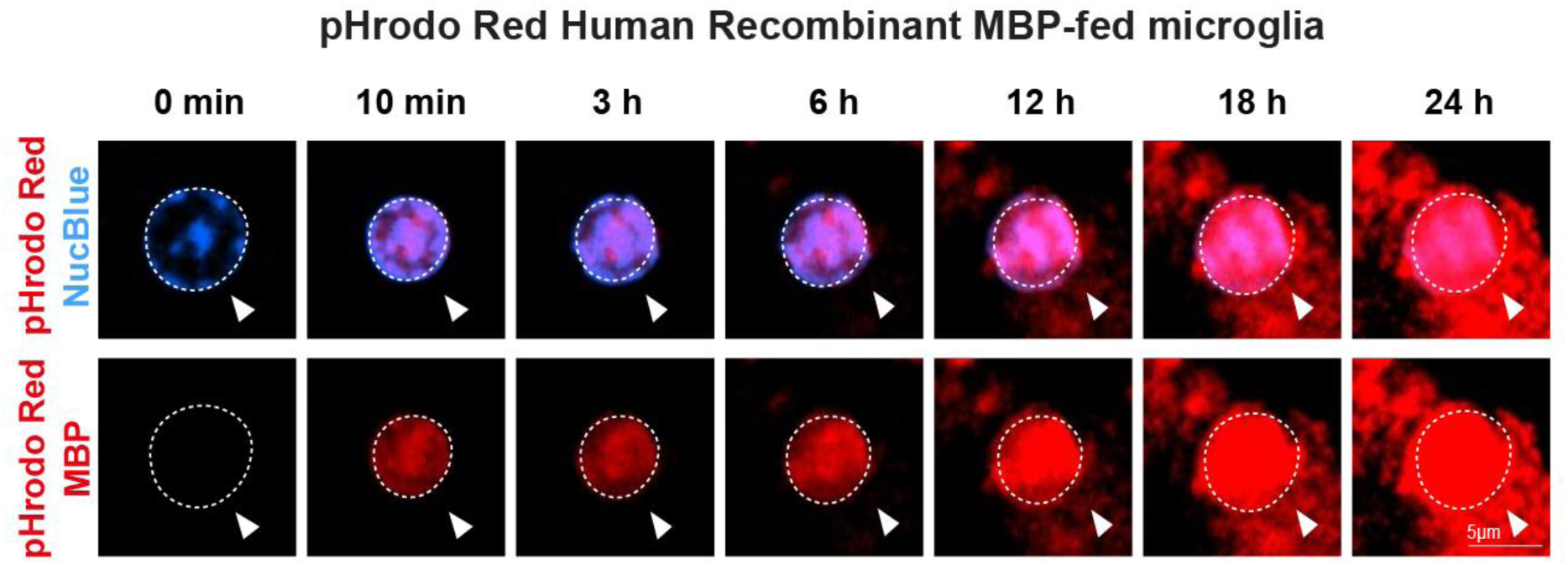
Further evidence of MBP localization to MG nucleus. Representative fluorescence micrographs of microglia exposed to pHrodo Red–labelled human recombinant MBP over the indicated time course (0 min to 24 h). Nuclei are counterstained with NucBlue^TM^ (blue). Dashed outlines indicate the cell boundaries and arrowheads mark regions of intracellular MBP accumulation. The upper row shows merged images, and the lower row shows the pHrodo Red channel alone. Progressive accumulation and increased pHrodo signal intensity are observed from early time points (10 min) through 24 h, including increased nuclear signal at later time points (6–24 h). Scale bar, 5 µm.

**Extended Data Fig 5.**
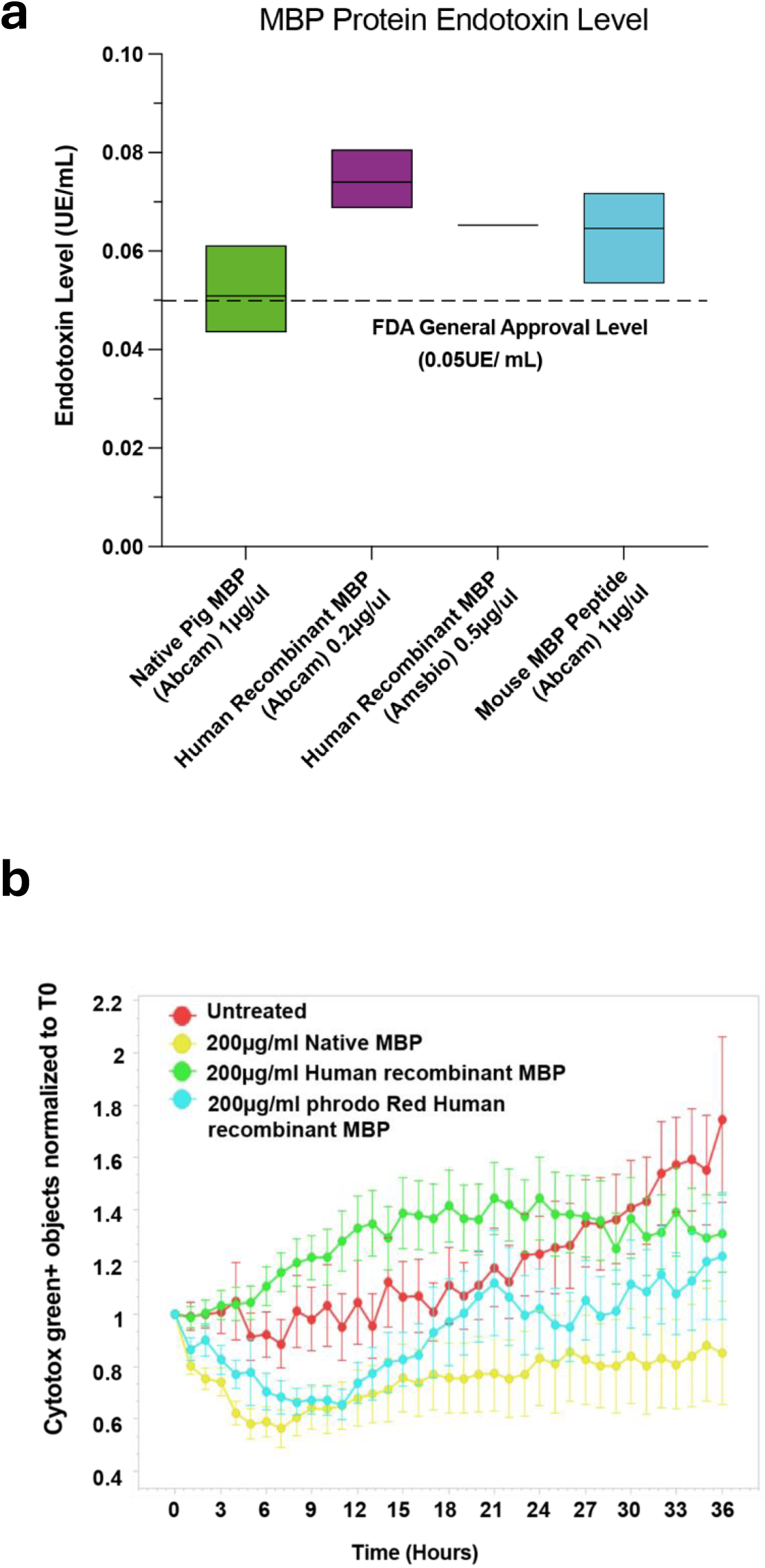
MBP variants used characterized for endotoxin levels. **a**, Quantification of endotoxin levels in myelin basic protein (MBP) preparations used in this study. Endotoxin levels were measured using the Pierce Chromogenic Endotoxin Quantification Kit (Thermo Fisher Scientific, A39552S). Data are presented as mean ± s.d. (*n* = 3). Native pig MBP (Abcam): 0.0509 ± 0.0095 EU ml⁻¹; recombinant human MBP (Abcam): 0.0740 ± 0.0063 EU ml⁻¹; recombinant human MBP (AMSBIO): 0.0653 EU ml⁻¹; mouse MBP peptide (Abcam): 0.0646 ± 0.0101 EU ml⁻¹. Dashed line indicates the general acceptable endotoxin threshold (0.05 EU ml⁻¹). **b**, Real-time quantification of cell death during longitudinal live-cell imaging. Data are presented as mean ± s.d. (*n* = 3). Cytotoxicity was assessed using the Incucyte® Cytotox Green assay (Sartorius, 4633), with the number of green objects (dead or dying cells) normalized to baseline (T₀). Conditions: untreated (red), native pig MBP (yellow), recombinant human MBP (green), and pHrodo Red–labelled recombinant human MBP (cyan), all at 200 µg ml⁻¹. MBP treatment does not induce detectable acute cytotoxicity over the imaging period.

**Extended Data Fig 6.**
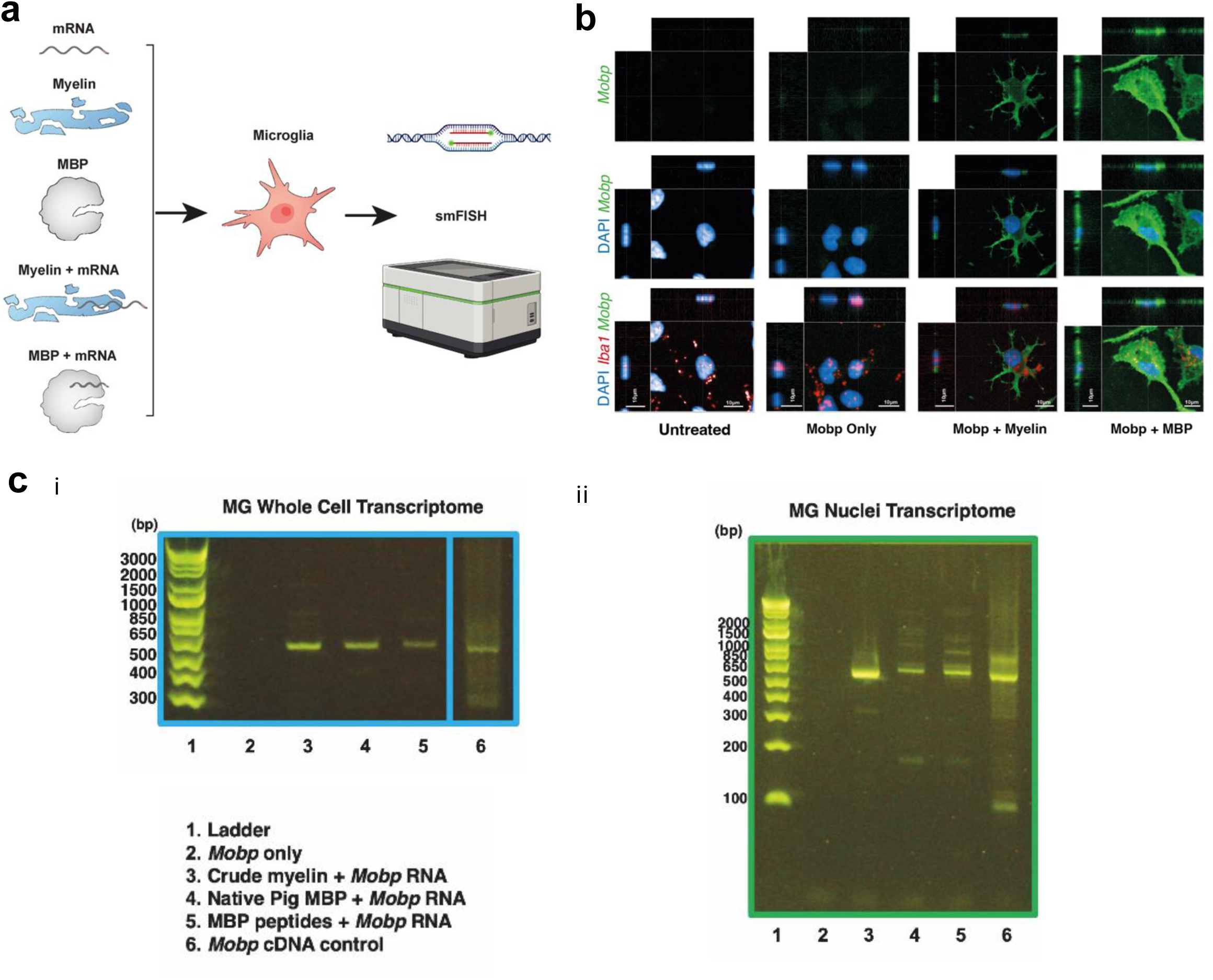
Nuclear retro-trafficked mRNA transcripts to the nucleus and preserved integrity. **a**, Schematic illustrating smFISH-based detection of MBP- or myelin-mediated trafficking of exogenous *Mobp* mRNA in microglia. Cells were exposed to myelin, MBP, *Mobp* mRNA alone, or MBP–mRNA complexes, followed by detection of intracellular transcripts using single-molecule fluorescence *in situ* hybridization (sm FISH). **b**, Representative smFISH images of microglia subjected to the indicated treatments (untreated, *Mobp* mRNA only, *Mobp* mRNA + myelin, or *Mobp* mRNA + MBP). *Mobp* mRNA signal (green) is detected by smFISH; microglia are labelled with IBA1 (red), and nuclei are counterstained with DAPI (blue). Scale bar, 10 µm. **c**, Agarose gel electrophoresis of RT–PCR products detecting *Mobp* transcripts. RNA was isolated from (i) whole-cell microglia (left) or (ii) microglial nuclear fractions (right) following the indicated treatments. Detection of *Mobp* signal in nuclear fractions indicates the presence of myelin-derived RNA following MBP-mediated uptake. Lanes: 1, molecular weight marker; 2, *Mobp* only; 3, crude myelin + *Mobp*; 4, MBP + *Mobp*; 5, MBP peptides + *Mobp*; 6, *Mobp* cDNA control.

**Extended Data Fig 7.**
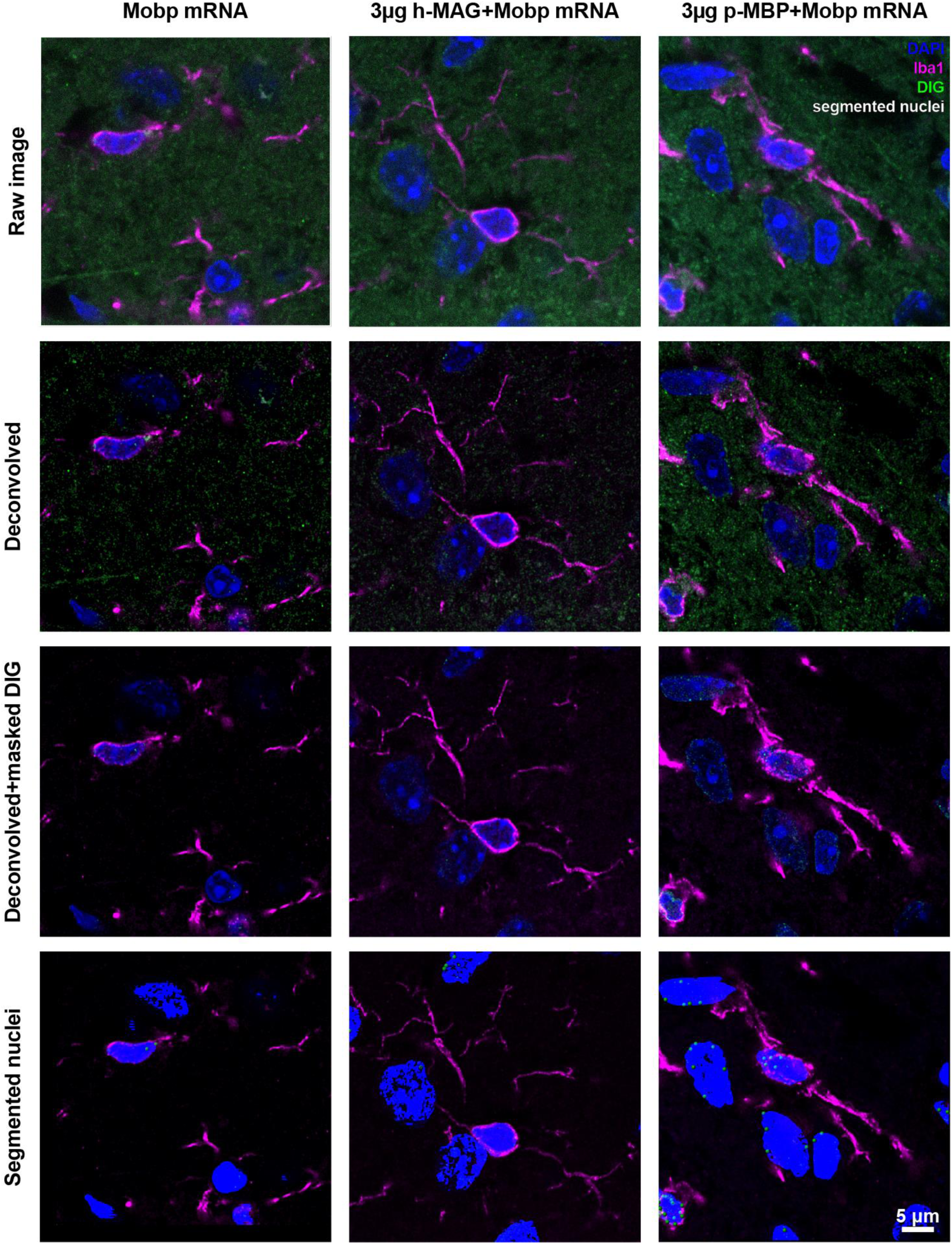
Further evidence for *in vivo* quantification of MBP-mediated RNA nuclear delivery in MG. Representative 0.5 μm thick Imaris-rendered micrographs of microglia cell bodies (Iba, magenta) from *Mobp* mRNA (left), *Mobp* mRNA + 3 µg h-MAG (middle) or *Mobp* mRNA + 3 µg p-MBP (right) injected animals. Micrographs depict a larger area of the examples used in Fig 6g, describing the image processing pipeline. Raw images were deconvolved using Huygens Professional (SVI) and imported to Imaris 10.2 (Bitplane). DAPI fluorescence (blue) was used to segment nuclei into surfaces and to generate a mask for nuclear DIG (green). Then, using the Cells applet, nuclei and DIG particles were segmented and quantified. Iba1 (magenta) was used to manually identify microglia. Segmented nuclei can be seen (blue) containing DIG particles, rendered as spheres (green). Scale bar: 10 μm.

## Notes

### Competing Interest Statement

The authors have declared no competing interest.

### Summary of Updates

In Version 5, we have fixed figure panel misalignment.

